# Functional and genetic determinants of mutation rate variability in regulatory elements of cancer genomes

**DOI:** 10.1101/2020.07.29.226373

**Authors:** Christian A. Lee, Diala Abd-Rabbo, Jüri Reimand

## Abstract

**Background:** Cancer genomes are shaped by mutational processes with complex spatial variation at multiple scales. Entire classes of regulatory elements are affected by local variations in mutation frequency. However, the underlying mutational mechanisms with functional and genetic determinants remain poorly understood.

**Results:** We characterised the mutational landscape of 1.3 million gene regulatory and chromatin architectural elements in 2,419 whole cancer genomes with transcriptional and pathway activity, functional conservation and recurrent driver events. We developed RM2, a statistical model that quantifies mutational enrichment or depletion in classes of genomic elements through genetic, trinucleotide and megabase-scale effects. We report a map of localised mutational processes affecting CTCF binding sites, transcription start sites (TSS) and tissue-specific open-chromatin regions. We show that increased mutational frequency in TSSs correlates with mRNA abundance in most cancer types, while open-chromatin regions are generally enriched in mutations. We identified ∼10,000 CTCF binding sites with core DNA motifs and constitutive binding in 66 cell types that represent focal points of local mutagenesis. We detected site-specific mutational signatures, such as SBS40 in open-chromatin regions in prostate cancer and SBS17b in CTCF binding sites in gastrointestinal cancers. We also proposed candidate drivers of localised mutagenesis: *BRAF* mutations associate with mutational enrichments at CTCF binding sites in melanoma, and *ARID1A* mutations with TSS-specific mutations in pancreatic cancer.

**Conclusions:** Our method and catalogue of localised mutational processes provide novel perspectives to cancer genome evolution, mutagenesis, DNA repair and driver discovery. Functional and genetic correlates of localised mutagenesis provide mechanistic hypotheses for future studies.

## Introduction

Genomes accumulate somatic mutations through exposure to exogenous and endogenous mutagens. Subsets of these mutations confer cells select proliferative advantages and drive oncogenesis, while most mutations are functionally neutral passengers ^1,2^. The discovery and validation of driver mutations is a major focus of cancer genomics research ^3-5^. However, the genome-wide landscape of passenger mutations is also instrumental to our understanding of oncogenesis and tumor evolution ^6,7^. Somatic mutation frequencies show complex genomic variation at multiple resolutions ^8^. In megabase-scale genomic windows, variations in mutation frequencies are associated with transcriptional activity, chromatin state and DNA replication, as late-replicating and non-transcribed regions are often more mutated than regions of early replication and highly expressed genes ^9-12^. At the base pair resolution, certain trinucleotides are preferentially mutated through processes of carcinogen exposures, defective DNA repair pathways, and aberrant DNA replication ^13-15^. For example, mutational signatures detected in metastatic tumors are informative of the treatment history of patients ^16,17^. In concert, these large-scale and nucleotide-level variations contribute to tumor heterogeneity and leave a footprint of tumor evolution and its cell of origin ^12,18,19^.

Complex variation in mutation frequencies is also apparent across intermediate genomic resolutions spanning hundreds to thousands of nucleotides. This encapsulates diverse functional genomic elements such as exons, transcription factor binding sites (TFBS) and chromatin architectural elements ^8,20^. DNA bound by nucleosomes and transcription factors (TFs) show increased mutation frequencies in cancer genomes ^21-23^. Active promoters in melanoma are enriched in UV-induced C>T somatic mutations resulting from differential activity of nucleotide excision repair influenced by DNA-binding of regulatory proteins ^24,25^. Likewise, DNA-binding sites of the master transcriptional regulator and chromatin architectural protein CTCF (CCCTC-binding factor) are enriched in somatic mutations in multiple cancer types ^21,26-28^. In contrast, certain genomic elements, such as chromatin-accessible regulatory regions ^29^ and protein-coding exons ^30^, have been shown to harbor relatively fewer mutations due to increased DNA repair activity. While the majority of such non-coding mutations tend to represent functionally neutral passengers, some regulatory elements at the high end of the mutation frequency spectrum may undergo positive selection due to their effects on cancer phenotypes. For example, the mutation hotspot in the *TERT* promoter creates a TFBS of the ETS TF family that leads to constitutive activation of *TERT* and enables replicative immortality of cancer cells ^31-33^. Recent studies have catalogued candidate non-coding driver elements in gene regulatory and chromatin architectural regions of the cancer genome with functional validations of novel elements ^34-36^ and shown that non-coding mutations converge on molecular pathways and regulatory networks involved in oncogenesis ^37,38^. Thus, we need to characterise localised mutational processes to deconvolute the role of carcinogens and endogenous mutational processes and the effects of positive selection in the non-coding genome. However, few dedicated computational methods exist to analyse variation in mutation frequencies at this resolution. As a result, there is a lack of large-scale analyses of the local mutation landscape in pan-cancer WGS datasets, leaving patterns of mutational enrichment and depletion undetected, and the genetic and environmental determinants poorly understood.

Here we developed a new statistical framework that quantifies the activity of mutational processes and signatures on specific classes of non-coding elements of the cancer genome. Our model considers local sequence context, megabase-level somatic mutation rates and genetic covariates to control for variation at the trinucleotide and megabase resolution while isolating site-level effects. We performed a systematic analysis of local mutation frequency variation in three classes of gene-regulatory and chromatin architectural genomic elements across 2,419 whole cancer genomes of the ICGC/TCGA Pan-cancer Analysis of Whole Genomes (PCAWG) project ^3^. We found a pervasive mutational enrichment at these functional non-coding elements that was characterised by specific mutational signatures and transcriptional and pathway-level activities in select cancer types. We detected statistical interactions of local mutagenesis and recurrent genomic alterations that suggest potential genetic mechanisms driving the underlying mutational processes. Our computational framework and systematic analysis reveal the diversity of mutational processes in functional non-coding elements of the cancer genome and their roles in somatic genome evolution, cancer phenotypes and molecular heterogeneity.

## Results

### A statistical framework for quantifying localised mutagenesis in cancer genomes

We implemented a statistical model, Regression Models for Local Mutations (RM2), to quantify the local activity of mutational processes in functional elements (*i*.*e*., sites) of whole cancer genomes, each spanning tens to hundreds of nucleotides (**Figure 1A**). The model considers a genome-wide set of elements, such as TFBSs isolated from chromatin immunoprecipitation with DNA sequencing (ChIP-seq), detected in thousands to hundreds of thousands of loci. The model uses negative binomial regression to evaluate whether the genomic elements of interest are collectively subject to a different mutation frequency compared to control sequences upstream and downstream of these elements. Somatic single nucleotide variants (SNVs) and small insertions-deletions (indels) were analysed; however, the model can be extended to rare germline variation and other classes of variants such as structural variant breakpoints. The model considers four types of information to evaluate local mutation rates: 1) nucleotide sequence content of genomic elements and control sequences representing the potential space for mutagenesis, grouped by 96 trinucleotide contexts and one indel context (*nPosits*), 2) the counts of observed somatic mutations in the cohort of tumors (*nMut*) in genomic elements and control sequences also grouped by 96 trinucleotide signatures and one indel signature required to derive mutation frequencies (*triNucMut*), 3) megabase-scale background mutation rates of elements computed across the cohort of tumors (*MbpRate*) to account for large-scale mutation correlates, such as transcription and chromatin state, and 4) an optional binary cofactor (*coFac*) to stratify tumors based on their genetic makeup (*e*.*g*., presence of a driver mutation) or clinical information (*e*.*g*., tumor subtype or stage). Genomic elements and flanking control regions are pooled into equally sized bins based on their megabase-scale mutation frequencies (ten bins by default). Elements and flanking control sequences are distinguished using the binary cofactor *isSite*. The full model is written as follows:

**Figure 1.**
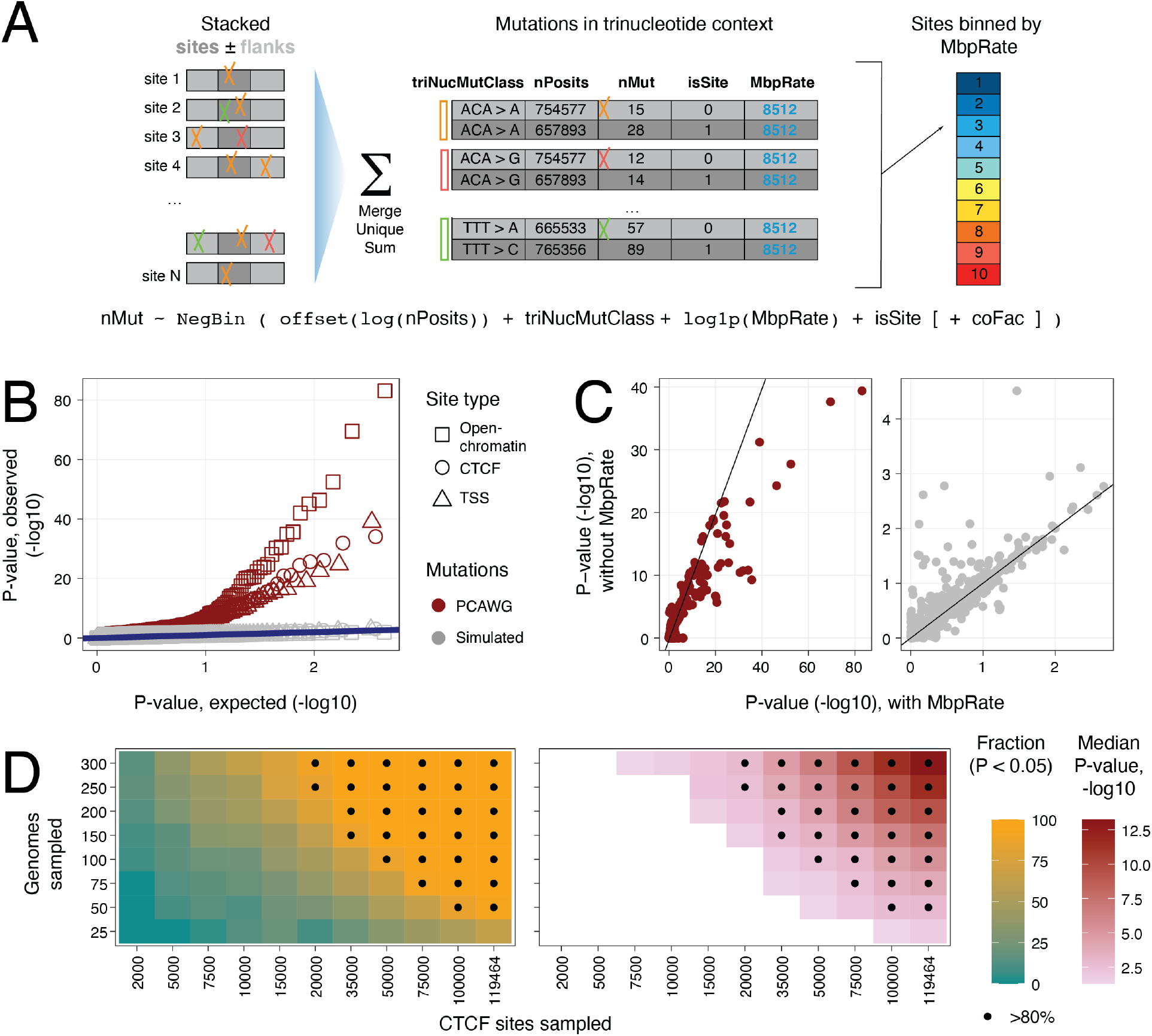
Characterising local mutational processes with RM2. **A**. Method overview. RM2 studies a set of genomic elements (*i.e*., sites) and somatic mutations in cancer genomes using a negative binomial regression model. Sites of constant genomic width (dark gray) and two control flanking sequences (light gray) are used *(isSite)*. Sites and flanks are collapsed into unique nucleotides and grouped to ten bins using their megabase-scale mutation frequency *(MbpRate)*. Mutations in sites and flanks *(nMuf)* are grouped by trinucleotide type *(triNucMut&ass)*. Trinucleotide content corresponding to the potential genomic space for mutations is used as model offset *(n Posits)*. Log-likelihood tests are used to compare the mutation frequencies in sites and flanking regions by removing the model cofactor *isSite*. The optional cofactor *coFac* enables interaction analysis of genetic and clinical variables. **B**. QQ-plot shows the observed and expected P-values of true and simulated mutations from PCAWG. No significant signals were identified in simulated data (FDR < 0.05), indicating that our method is well-calibrated. **C**. Comparison of model performance with and without *MbpRate* covariate. Analysis of true (left) and simulated mutations (right) shows the advantage of modelling megabase-scale mutation frequency. **D**. Power analysis of RM2 using down-sampling of CTCF binding sites and liver cancer genomes. Fraction of significant results (left) and median P-value (right) are shown. Panels B-C include total mutations, strand- and signature-specific mutations as in Figure 2. Only total mutations were included for analyses in C-D.

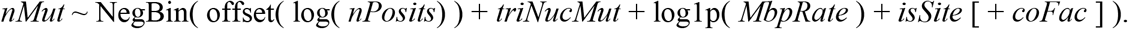

To determine whether the mutation frequencies of genomic elements differ from flanking sequences given trinucleotide-level and megabase-scale covariates, we evaluate the significance of the cofactor *isSite* using a likelihood-ratio test. Significant and positive coefficients of this cofactor indicate increased mutation rates in genomic elements relative to flanking controls, while negative coefficients indicate a depletion of mutations. Similarly, we can discover potential genetic or clinical interactions with localized activity of mutational processes. Given a binary subgroup classification of tumors (*coFac*), we evaluate the significance of its interaction with local mutation rates (*isSite:coFac*). Positive coefficients of the interaction indicate that the mutation rates in a clinical or genetic tumor subgroup are elevated when accounting for the overall differences of the subgroups. We also extend the analysis to classes of mutations, such as those of COSMIC mutational signatures, by allowing only specific classes to be included in the mutation counts (*nMut*). We evaluated the performance of our method using simulated datasets, power analysis and parameter variations as described below (**Figure 1B-D**).

### Comprehensive map of mutational processes in gene-regulatory and chromatin architectural elements of cancer genomes

To study localised mutation frequencies in gene-regulatory and chromatin architectural elements, we used the pan-cancer dataset of 2,514 whole cancer genomes of the PCAWG project ^3^ with high-confidence SNVs and indels. We individually analysed 25/35 cancer types with at least 25 samples, as well as the pan-cancer set of all 35 cancer types (**Supplementary Figure 1A**). Hypermutated tumors (69 or 2.7%) were excluded to avoid confounding effects. To perform a conservative analysis, we excluded a few tumors as outliers (33 or 1.4%) where even single-sample RM2 analysis revealed highly significant mutational enrichments (*FDR* < 0.001) (**Supplementary Figure 1B**). This ensured that our findings were shared across most tumor genomes and did not represent isolated trends representative of few tumors. The final analysis considered 2,419 cancer genomes of 35 cancer types with 22.7 million mutations including 1.62 million indels. In addition to all mutations combined, we grouped the mutations by COSMIC mutational signatures of single base substitutions (SBS) inferred in the PCAWG study ^14^, reference and alternative nucleotide pairs, and DNA strand information. Indel mutations were pooled with SNVs and also analysed separately.

Three classes of genomic elements spanning 1,269,347 unique loci and 10% (337.0 Mbps) of the human genome were analysed. These included 119,464 CTCF binding sites conserved in at least two cell lines in the ENCODE project ^39^, 37,309 transcription start sites (TSS) of protein-coding genes from the Ensembl database (GRCh37), and 1,193,391 unique open-chromatin sites. Open-chromatin sites were analysed in tissue-specific subsets, ranging from 43,000 to 500,000 sites per cancer type (**Supplementary Table 1A**). For most cancer types (17/25), matching open-chromatin sites were adapted from the ATAC-seq profiles of primary tumors of the Cancer Genome Atlas (TCGA) project ^40^. The pan-cancer analysis used 500,183 pan-cancer sites from TCGA. For eight cancer types, DNAse-seq profiles of the closest relevant normal tissues of the Roadmap Epigenomics project ^41^ were used. Open-chromatin sites were filtered to exclude TSSs and CTCF binding sites to allow direct comparison and reduce confounding effects of the three site classes.

The analysis revealed a landscape of localised mutational processes in gene-regulatory and chromatin architectural elements of cancer genomes (**Figure 2A**). We found 307 significant differences in mutation frequencies in the three classes of genomic elements (RM2, FDR < 0.05). These affected 21 cancer types and included 23 unique mutational signatures. The majority of findings (281) indicated mutational enrichments at the sites compared to adjacent flanking regions, while a few reduced mutation frequencies were also observed (26 or 8%). The strongest cumulative signals were found in open-chromatin sites in prostate, liver and breast cancers and CTCF binding sites in liver and esophageal cancers.

**Figure 2.**
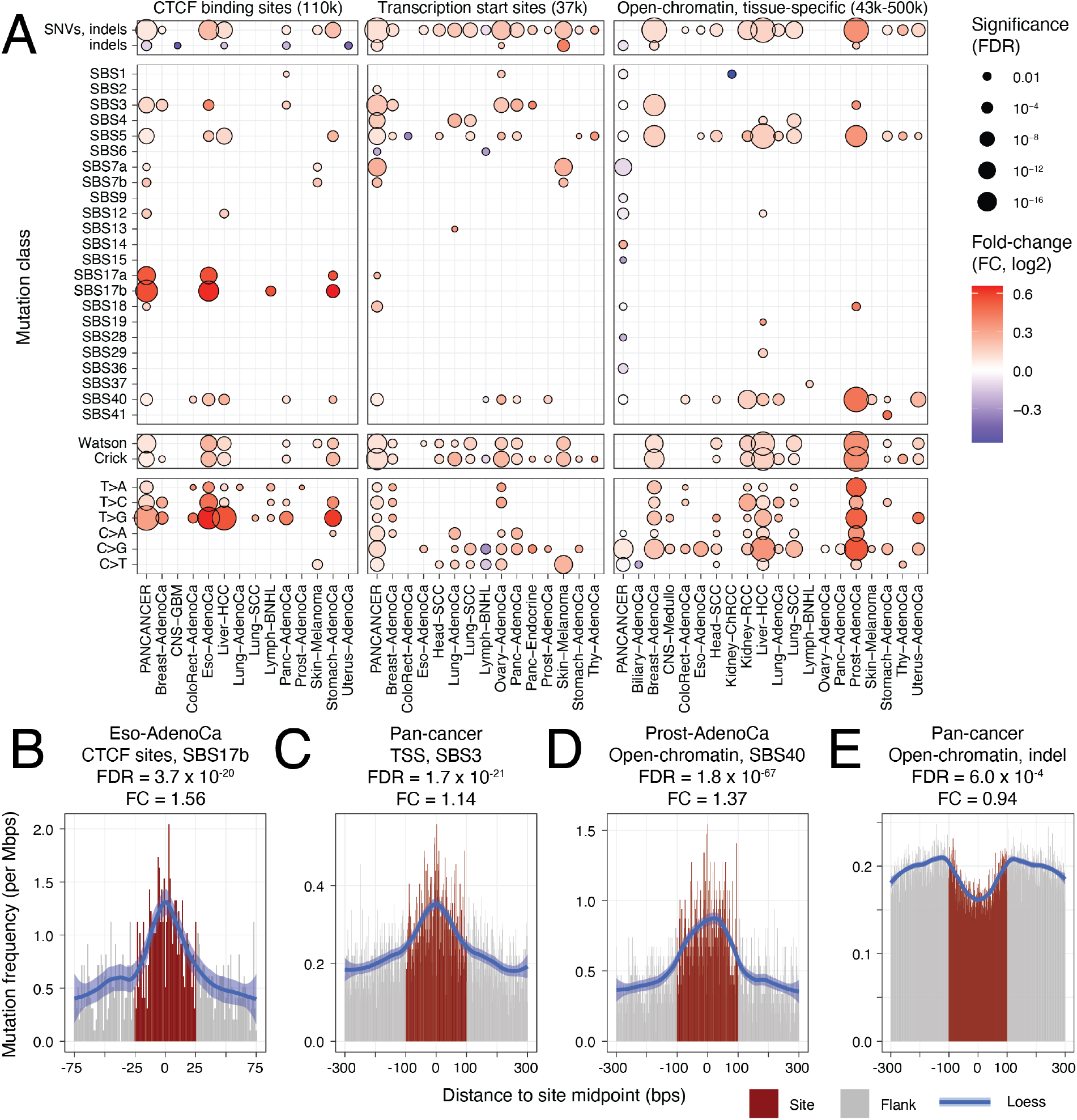
Comprehensive map of mutational processes in gene-regulatory and chromatin architectural elements of cancer genomes. **A**. Comparison of mutation frequencies in DNA-binding sites of the CTCF chromatin architectural factor (left), transcription start sites (TSS) and cancer-specific open-chromatin sites in 2,419 whole cancer genomes *(FDR* < 0.05). Total mutations (SNVs, indels) and mutations grouped by single base substitution (SBS) signatures, substitution types and DNA strand were analysed. Open-chromatin sites were filtered to exclude TSSs and CTCF sites. B-E. Examples of localised mutation frequencies and signatures: **B**. Enrichment SBS17b in CTCF binding sites in esophageal adenocarcinoma, **C**. Pan-cancer enrichment of SBS3 mutations in TSSs. D. Enrichment of SBS40 mutations in open-chromatin sites in prostate adenocarcinoma. E. Pan-cancer depletion of indel mutations in open-chromatin sites.

We first focused on the mutational profiles of CTCF DNA-binding sites. Mutational enrichments in CTCF binding sites were the strongest in liver hepatocellular carcinoma (RM2 *FDR* = 1.22 × 10^−12^, fold-change (FC) = 1.08), esophageal adenocarcinoma (*FDR* = 6.0 × 10^−20^, FC = 1.17) and stomach adenocarcinoma (*FDR* = 6.2 × 10^−11^, FC = 1.16), and the pan-cancer cohort. Smaller enrichments were detected in melanoma, pancreatic and breast cancer (*FDR* ≤ 0.02). Subgroups of mutations revealed further signals. Strong enrichments of thymine base substitutions (T>G, T>C, T>A) were found in nine cancer types (*e*.*g*., T>G in Liver-HCC, *FDR* = 1.3 × 10^−30^, FC = 1.46), while cytosine base substitutions were not enriched. We then asked whether CTCF binding sites were characterised by COSMIC SBS signatures. The strongest site-specific enrichments were found for SBS17 in esophageal and stomach cancer and Non-Hodgkin’s lymphoma (SBS17b in esophageal cancer: *FDR* = 3.7 × 10^−20^, FC = 1.56) (**Figure 2B**). Interestingly, liver and other cancer types with overall frequent CTCF binding site mutations did not show a site-specific enrichment of SBS17. The aetiology of SBS17b is unknown, however it has been linked to acid reflux and oxidative damage to DNA in gastro-esophageal cancers ^42,43^, and a similar mutational signature found in metastatic tumors has been associated with the effects of nucleoside metabolic inhibitor chemotherapies capecitabine and 5-FU ^16^. Our analysis suggests that effects of these mutagens may be especially active at insulator and chromatin architectural elements bound by CTCF in tissues of the digestive system. These results confirm earlier reports of elevated mutation rates in CTCF DNA-binding sites ^21,27,28^ in a large and diverse dataset of whole cancer genomes, validating our computational model and refining the annotation of the mutational processes associated with CTCF binding sites.

TSS of protein-coding genes were significantly enriched in mutations in the pan-cancer cohort (*FDR* = 1.7 × 10^−37^, FC = 1.07) and in cohorts of 13/25 cancer types, most prominently in melanoma (*FDR* = 3.6 × 10^−14^, FC = 1.15), breast, head, lung, ovary and pancreatic cancers (*FDR* ≤ 10^−5^). Enrichments of cytosine base substitutions were found in melanoma (C>T, *FDR* = 1.4 × 10^−15^, FC = 1.19) as well as lung, ovarian and head and neck cancers. In contrast to CTCF binding sites, thymine base substitution frequencies were not elevated. Mutational signature analysis highlighted an elevated activity of the aging-associated signature SBS5 in the pancancer cohort (*FDR* = 2.4 × 10^−13^, FC = 1.07) and in eight cancer types. Signature SBS3, associated with defects of homologous recombination-based DNA damage repair, was enriched at TSSs in the pan-cancer cohort (*FDR* = 1.7 × 10^−21^, FC = 1.14) (**Figure 2C**) as well as breast, pancreatic and ovarian cancers (*FDR* ≤ 10^−5^). Spontaneous formation of endogenous DNA double strand breaks at promoters has been associated with the pause and release of RNA polymerase II and linked to chromosomal translocations in cancer ^44^, suggesting a mechanism of this TSS-specific mutagenesis in cancer genomes. TSSs were enriched in carcinogen-driven mutational signatures, such as the ultraviolet light signature in melanoma (SBS7a: *FDR* = 1.1 × 10^−14^, FC = 1.21) and the tobacco signature in two lung cancer cohorts (SBS4: *FDR* ≤ 10^−5^). These match the major mutagens and exposures of those cancer types, indicating an overall increased vulnerability of TSSs to mutational processes and carcinogens. This analysis extends previous reports of increased mutation frequencies in promoters in melanoma ^24,25^ to additional cancer types, demonstrating that TSS-specific mutational processes are widely active in cancer genomes.

Tissue-specific open-chromatin regions were also enriched in mutations in 12/25 cancer types, with the strongest signals in prostate cancer (FDR = 9.8 × 10^−81^, FC = 1.29) as well as liver, lung, breast and kidney cancer (FDR ≤ 10^−14^) whereas C>G mutations were more frequently observed compared to other base substitutions. Mutational signature analysis revealed the enrichment of SBS40 mutations in open-chromatin regions in nine cancer types and the pan-cancer cohort, with the strongest signal apparent in prostate cancer (FDR = 1.8 × 10^−67^, FC = 1.33) (**Figure 2D**). The aging-associated signature SBS5 was enriched in open-chromatin sites in eleven cancer types, especially in liver, prostate, breast and lung cancer (FDR ≤ 10^−9^). In contrast to strong signals in individual cancer types, the analysis of pan-cancer open-chromatin sites and mutations revealed only a trend of elevated mutation frequencies at open-chromatin sites (FDR = 0.056, FC = 1.01). The pan-cancer sites were depleted of indel mutations (*FDR* = 6.0 × 10^−4^, FC = 0.94) (**Figure 2E**) and several less-frequent mutational signatures, while those effects were not observed in individual cancer types. The limited findings of the pan-cancer analysis emphasize the importance of integrating open-chromatin profiles of matched cancer types to study the interactions of chromatin and mutagenesis. Overall, open-chromatin regions of individual cancer types are enriched in mutations through specific mutational processes acting at these local scales.

We used the generated map to benchmark our model and compared it to a PCAWG dataset of simulated variant calls designed to approximate neutral genome evolution ^4^. First, analysis of simulated data did not reveal any significant differences in mutation frequencies in the three classes of elements (all *FDR* > 0.05). Quantile-quantile analysis of *P*-values confirmed that the model is well calibrated for true and simulated mutations (**Figure 1B**). Second, we compared the models with and without accounting for the megabase-scale mutation frequency covariate (MbpRate) and found that including the information led to higher significance in the true mutation dataset and lower significance in the simulated mutation dataset (**Figure 1C**). Third, we evaluated the statistical power of our model by analyzing total mutations in down-sampled subsets of liver cancer genomes and CTCF binding sites (**Figure 1D**). For example, the mutational enrichment in CTCF sites was detectable 80% of the time when sampling 75 genomes and 75,000 CTCF binding sites. Fourth, we varied the parameter corresponding to the normalised width of genomic elements for the three classes (**Supplementary Figure 2A**). Site-specific differences in mutation frequencies were robustly detected for multiple genomic widths of sites. However, mutational enrichments in site bound by CTCF were generally narrower (50 bps) compared to TSS and open-chromatin sites (200 bps), indicating differences in mutational processes. In summary, our method provides a versatile and well-calibrated framework for analysing localised mutational processes in cancer genomes.

### Mutational enrichment at transcription start sites associates with mRNA abundance of target genes and diverse pathways

We asked whether the increased mutation frequency at TSSs and open-chromatin sites correlated with transcription of target genes. We used matched RNA-seq data available for 20 cancer types in PCAWG ^45^ to quantify the tissue-specific activity of regulatory regions. Protein-coding genes were distributed into five equally-sized bins based on their median mRNA abundance in each cancer type such that the first bin included silent genes and the fifth bin included a wide range of highly transcribed genes. Open-chromatin sites were assigned to genes based on their location in gene bodies and promoters, as well as consensus long-range chromatin interactions in promoter-capture Hi-C experiments ^46^. This provided a tissue-specific map of TSSs and open-chromatin sites with transcript abundance as a measure of site activity.

Higher mutation frequencies in TSSs strongly associated with tissue-specific mRNA abundance of target genes (**Figure 3A**). In the fifth bin of highly transcribed genes, TSSs were consistently enriched in mutations in 12/20 cancer types (*FDR* < 0.05) and the pan-cancer cohort (*FDR* = 6.2 × 10^−38^, FC = 1.15) (**Supplementary Table 1C**). The strongest associations of transcriptional activity and TSS-specific mutagenesis were found in melanoma, ovarian, lung, pancreatic and breast cancer (*FDR* ≤ 10^−4^, FC ≥ 1.15) (**Figure 3B**,**C**). The TSSs of genes with intermediate transcript abundance (40-80 percentile) were also enriched in mutations in seven cancer types. In contrast, TSSs of silent and low-abundance genes were not differentially mutated compared to flanking controls. Interestingly, the overall mutation frequency was higher in and around silent TSSs, suggesting that closed chromatin is exposed to lower levels of DNA repair. Of note, the top gene bin was highly variable in mRNA abundance (*e*.*g*., range 11—7100 FPKM-UQ in the pan-cancer cohort) **(Figure 3C)**. Therefore, the mutational enrichment in the top gene bin may be partially contributed to by genes with very high expression. TSS bins with higher transcript abundance were also enriched in mutational signatures commonly observed in their respective cancer types (**Supplementary Figure 3A**). This analysis highlights a pan-cancer mutational process localised in TSSs that appears to be driven by transcriptional activity.

**Figure 3.**
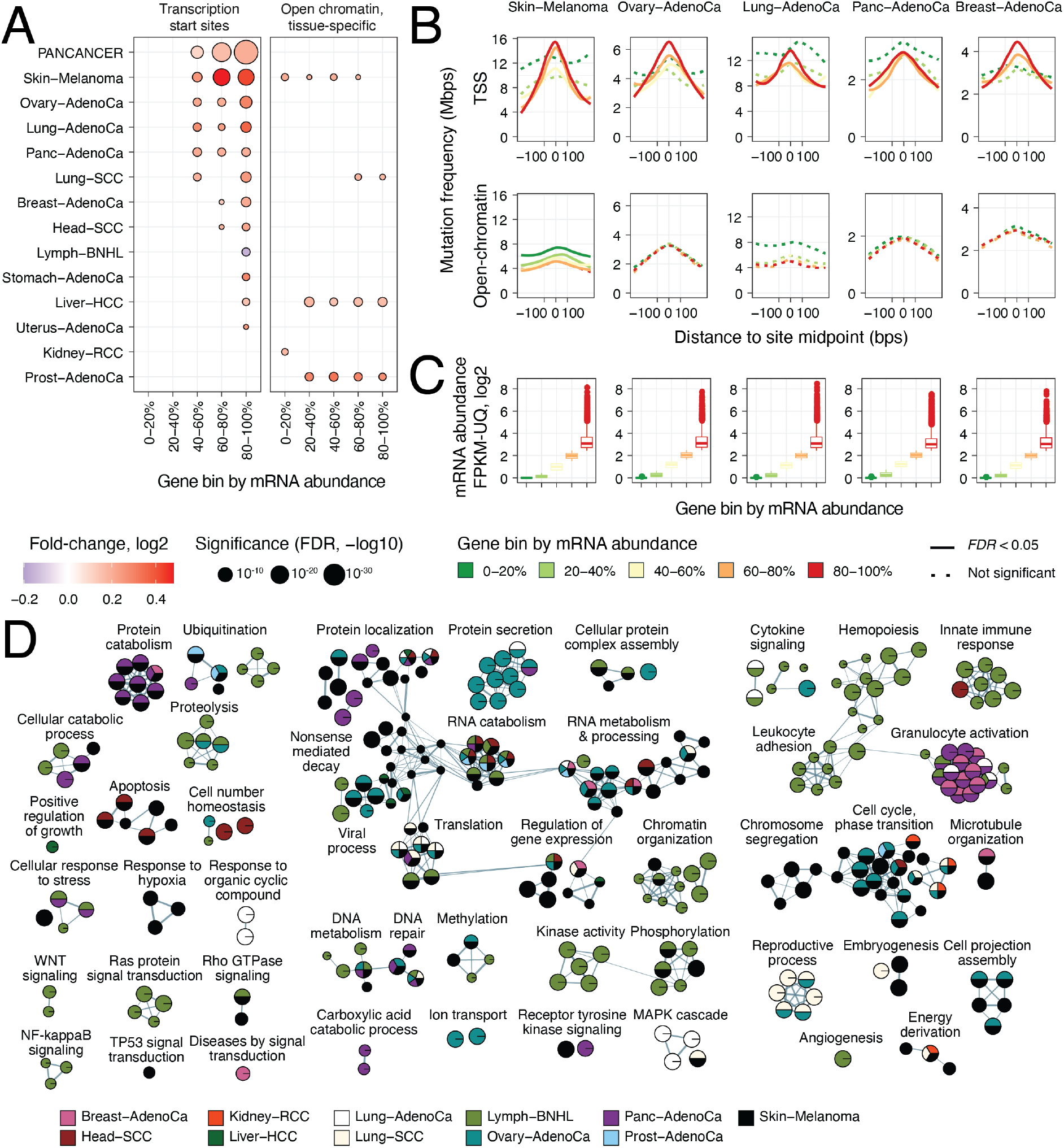
Mutational enrichment at transcription start sites associates with mRNA abundance of target genes and diverse pathways. **A**. Comparison of mutation frequencies in TSSs and tissue-specific open-chromatin sites grouped by mRNA abundance of target genes in matching tumors (FDR < 0.05). Equal numbers of sites were sampled for an unbiased comparison. **B**. Examples of cancer types with strong transcription-associated mutagenesis in TSSs (top) compared to open-chromatin sites (bottom). Mutation frequencies are shown on the Y-axis with loess smoothing. **C**. Median mRNA abundance of genes in the five bins. **D**. Enrichment map of pathways and processes with frequent mutations at TSSs (FDR < 0.05). Nodes represent pathways and processes that are connected with edges if these include many common genes.

Compared to TSSs, mutation frequencies in open-chromatin sites were attenuated and did not associate with transcript abundance of target genes (**Figure 3A,B**). Mutational enrichments in multiple gene bins were observed in melanoma, liver cancer and prostate cancer, however the effect sizes were smaller, and most cancer types showed no significant changes in local mutation frequencies. In this unbiased comparison, we included equally-sized bins of TSSs and open-chromatin sites using down-sampling, since open-chromatin sites considerably outnumbered TSSs. An extended analysis of all open-chromatin sites grouped by transcript abundance showed stronger mutational enrichments in most cancer types that is consistent with our analysis above (**Supplementary Figure 2B**), however, no associations with transcript abundance were apparent. This analysis suggests that transcription-associated local mutagenesis is more prominent in TSSs than in general regions of open chromatin.

To explore the functional associations of elevated mutation frequencies at TSSs, we performed a pathway enrichment analysis by adapting RM2 to gene sets of GO biological processes and Reactome molecular pathways ^47^. This allowed us to test our hypothesis that the TSSs enriched for mutations are concentrated in specific biological processes. We found 336 unique pathways and processes with pronounced enrichments of mutations at TSSs in eleven cancer types (*FDR* < 0.05) (**Figure 3D**) (**Supplementary Table 1D**). Half of the pathways were found in at least the melanoma cohort (51%). However, one third of the pathways were found in two or more cancer types, indicating that the pathway associations of elevated mutagenesis at TSSs often apply more generally to multiple cancer types. Translation, ribosome biogenesis and RNA processing were among the largest groups of pathways found. This is expected as the translational machinery is ubiquitously active in proliferating cells and includes many highly expressed genes. Processes related to gene regulation and chromatin organization were also detected. Besides these general housekeeping processes, cancer-related processes and pathways were also enriched in TSS mutations. For example, mitotic cell cycle, apoptosis, DNA repair, angiogenesis, developmental and immune response processes, as well as druggable signalling pathways (*e*.*g*., MAPK, Wnt, Notch) were identified in multiple cancer types. The pathway analysis supports our findings of frequent TSS mutations associated with increased transcription. It also highlights a variety of core cellular processes and cancer pathways where mutations accumulate in core promoters across multiple cancer types and may have functional consequences.

### Mutational enrichment at CTCF binding sites is associated with constitutive DNA-binding and core sequence motifs

We tested whether the elevated mutation frequencies in CTCF binding sites were associated with functional site characteristics. We used the extent of conservation of DNA-binding across 70 cell lines as a proxy of site activity. We grouped 162,209 unique CTCF binding sites catalogued in ENCODE into five equal bins such that the first bin contained sites observed in a single cell line while the fifth bin included constitutively bound sites with the median site found in 67/70 cell lines (**Figure 4A**). Doing so allowed us to test mutation frequency and genomic associations at different levels of site activity. The constitutively bound CTCF binding sites were enriched in chromatin loop anchors ^48^ (29% observed *vs*. 12% expected, Fisher’s exact *P* < 10^−300^) and consensus core CTCF DNA-binding motifs (48% observed *vs*. 29% expected, *P* < 10^−300^) (**Figure 4B**). Together, these features imply that the constitutive bin of sites represents functionally important CTCF binding sites.

**Figure 4.**
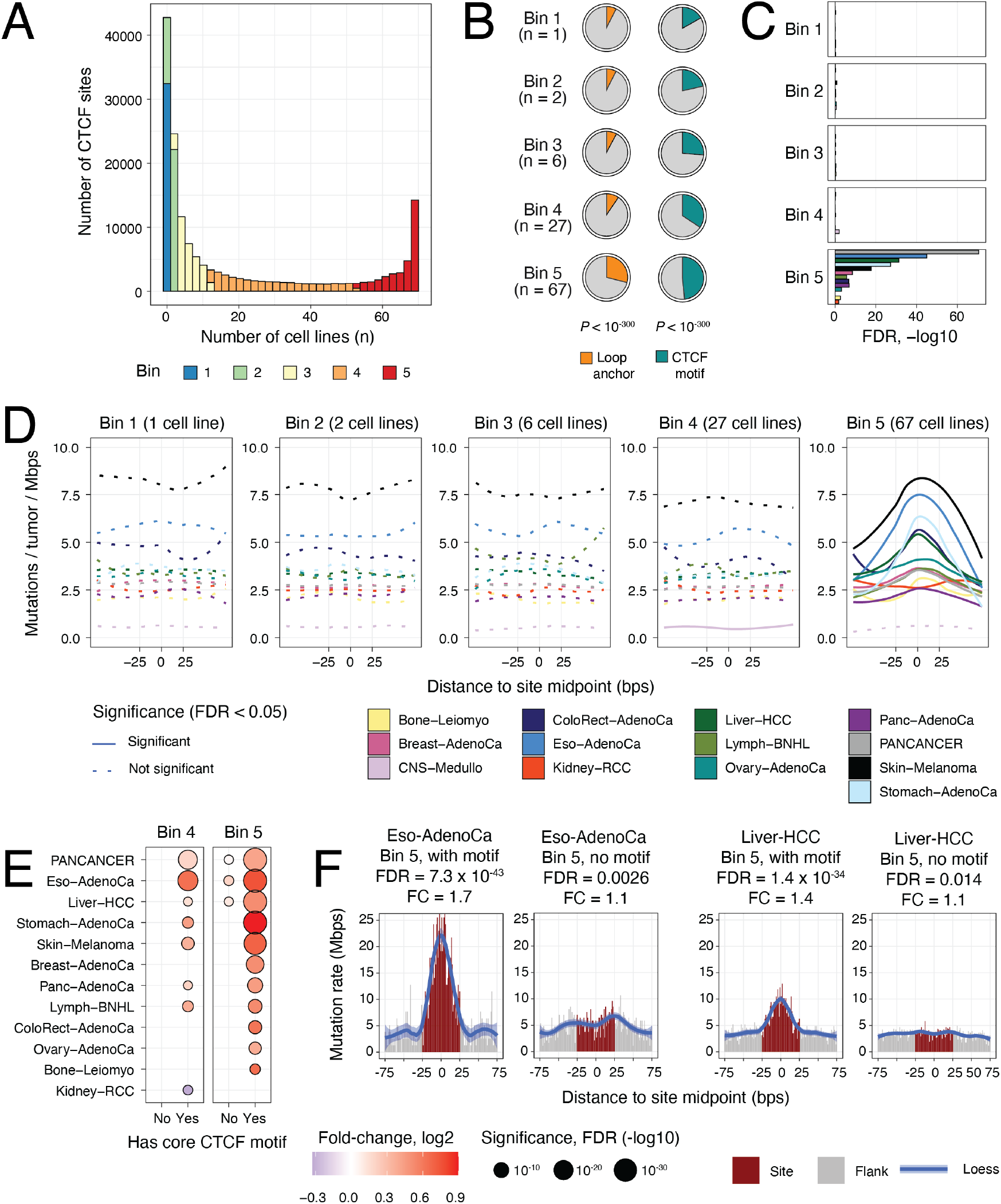
Localised mutational processes at constitutively active binding sites of CTCF. **A**. Histogram of CTCF binding sites with number of cell lines in ENCODE. Sites were grouped as five equal bins based on conservation across cell lines (colors). Bimodal distribution reveals a subset of sites detected in most or all cell types (in red). **B**. Pie charts show the proportion of CTCF sites in the five bins located at chromatin loop anchors (left) and that matched the core CTCF motif (right). P-values represent the enrichments in the 5th bin compared to all sites (Fisher’s exact test). The median numbers of cell lines per bin are shown in brackets. **C**. Significance of the localised mutational enrichments in the five bins of CTCF binding sites. FDR values of the RM2 analysis are shown on the X-axis. Colors correspond to cancer types. **D**. Local mutation frequency in the five bins of CTCF binding sites. Solid lines show statistically significant changes in mutation frequencies compared to flanking controls (RM2 FDR < 0.05). Loess curves were used for smoothing. **E**. Significance of mutational enrichment in highly conserved CTCF binding sites (bins 4-5) grouped by presence or absence of core DNA-binding motif of CTCF in the sites. **F**. Mutations in constitutively bound subsets in CTCF binding sites with and without the core CTCF DNA motifs. Esophageal cancer (left) and liver cancer (right) are compared.

The constitutive CTCF binding sites were characterised by elevated mutational frequencies in the pan-cancer cohort (*FDR* = 1.4 × 10^−71^, FC = 1.18) and in ten cancer types, especially stomach, liver and esophageal cancers as well as melanoma (*FDR* ≤ 10^−19^, FC ≥ 1.25) (**Figure 4C,D**) (**Supplementary Table 1E**). In contrast, all other bins of CTCF sites showed no significant enrichment of mutations (*FDR* > 0.05). Specifically, none of the cancer types showed mutational enrichments in the sites of the fourth bin which still represent a high level of conservation of CTFC binding (median 27/70 cell lines). Mutational signature analysis revealed over-represented signatures such as SBS3, SBS5, SBS7, SBS17 and SBS40 at these sites in various cancer types, however a common signature for constitutive CTCF binding was not apparent (**Supplementary Figure 3B**). In summary, this analysis emphasises the potential role of constitutive CTCF binding activity in local mutagenesis.

We then examined whether the presence of CTCF DNA-binding motifs in the ChIP-seq peaks played a role in localised mutagenesis. We categorised all the CTCF binding sites into two subsets based on the presence or absence of a consensus core CTCF DNA-binding motif and repeated the conservation-based grouping of the two subsets of sites. The localised mutational process at CTCF binding sites appears to be driven by the combination of constitutive binding and core motif presence (**Figure 4E**) (**Supplementary Table 1F**,**G**). The constitutively bound CTCF sites matching the DNA motifs were strongly enriched in mutations in ten cancer types and the pan-cancer cohort. In contrast, the constitutively-bound sites lacking the core DNA motif were not enriched. For example, esophageal cancer showed a strong mutational enrichment in the constitutively-bound CTCF sites with the core motif present (*FDR* = 7.3 × 10^−43^, FC = 1.70), while the signal was clearly attenuated in the constitutively-bound sites with no core motif (*FDR* = 0.0026, FC = 1.14) (**Figure 4F**). This was confirmed in liver cancer (bin 5 with motifs: *FDR* = 1.4 × 10^−34^, FC = 1.44; bin 5 without motifs: *FDR* = 0.014, FC = 1.08). Highly conserved sites with median conservation in 27/70 cell lines also showed motif-driven mutational enrichments. Strikingly, no signal of mutational enrichment was detected in any other bins of sites regardless of motif presence (**Supplementary Figure 4A**). The findings were confirmed in a tissue-specific analysis integrating breast and liver cancer mutations with CTCF binding sites of the cancer cell lines MCF-7 and HepG2, respectively (**Supplementary Figure 4B**,**C**).

Our analysis highlights a minority of CTCF binding sites (9,552 or 5.9%) with constitutive CTCF binding and core DNA motifs that are the primary target of a local mutational process in many cancer types. This analysis refines the established pattern of mutational enrichment in CTCF binding sites by defining a subset of frequently-mutated sites with specific functional properties, such as involvement in chromatin architectural and gene regulatory interactions ^48,49^. Our observations are consistent with previous findings of CTCF motifs in sites shared among a few cell lines that comprised a particularly strong mutational enrichment indicating higher site activity within cells ^50,51^. Conservation is a property of functionally integral CTCF binding sites, which upon disruption, can lead to changes in underlying chromatin architecture and gene regulation ^52^ and is associated with activation of proto-oncogenes ^26^. Therefore, the accumulation of mutations at those sites may have functional consequences in cancer.

### Recurrent driver mutations and copy-number alterations associate with localised mutagenesis

To find potential genetic mechanisms of localised mutagenesis, we tested whether the presence of specific features such as recurrent mutations in tumor genomes associated with higher mutation frequencies at gene-regulatory and architectural elements. To enable this analysis, we collected 37 driver genes with frequent SNVs and indels predicted using the ActiveDriverWGS method ^34^, 70 recurrent copy-number alterations (CNAs) detected in the PCAWG project using the GISTIC2 method ^53^, as well as two genome-wide measures of aneuploidy, whole-genome duplication (WGD) and percent genome altered (PGA) (**Supplementary Figure 5**). Integrative analysis with RM2 identified 41 significant interactions between genomic features and site-specific elevations in mutation frequencies in nine cancer types, including five driver genes (*ARID1A, BRAF, CTNNB1, HIST1H1C, SETD2*), 25 recurrently amplified loci and one deletion (RM2 *FDR* < 0.05, interaction *P* < 0.05) (**Figure 5A**) (**Supplementary Table 1G**).

**Figure 5.**
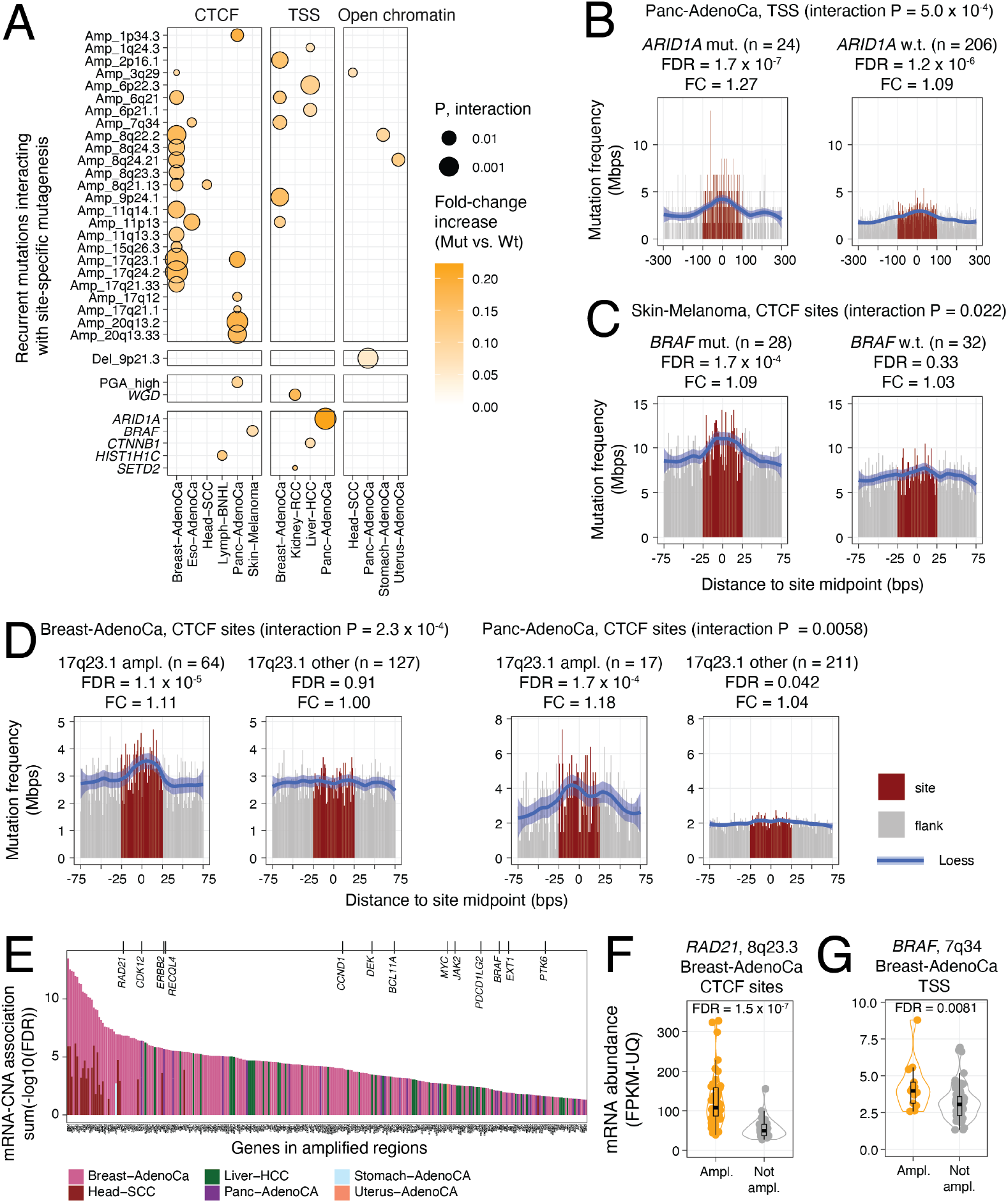
Recurrent driver mutations and copy-number alterations (CNA) associate with localised mutagenesis. **A**. Dotplot of the statistical interactions of recurrent mutations with increased mutation frequencies at sites (RM2 FDR < 0.05; interaction P < 0.05). Whole-genome duplication (WGD) and CNAburded (median-dichotomised percent genome altered, PGA) are also shown. **B-D**. Examples of increased mutational frequencies associating with recurrent mutations. Tumors with and without recurrent mutations are shown (left *vs*. right). **B**. *ARID1A* mutations in pancreatic cancer associate with enriched mutations at TSSs. **C**. *BRAF*mutations in melanoma associate with enriched mutations at CTCF binding sites. **D**. 17q23.1 amplifications associate with enriched mutations at CTCF binding sites in breast and pancreatic cancer. **E**. Copy-number amplified genes with amplification-driven increases in mRNA abundance. Known cancer genes are shown at the top. **F**. *RAD21* was up-regulated in the set of 8q23.3-amplified breast cancers associated with enriched mutations in CTCF binding sites. **G**. *BRAF was* up-regulated in the set of 7q34-amplified breast cancers associated with enriched mutations in TSSs.

*ARID1A* mutations in pancreatic adenocarcinoma had one of the strongest interactions with localised mutational enrichment (interaction *P* = 5.0 × 10^−4^) (**Figure 5B**). The 24 pancreatic cancer genomes with *ARID1A* mutations showed a significant increase in mutation frequency at TSSs (FDR = 1.7 × 10^−7^, FC = 1.27). In contrast, the majority of tumors (206) lacking *ARID1A* mutations showed a weaker, albeit significant, mutational enrichment at TSSs, likely explained by the improved statistical power of the larger set of samples analysed (FDR = 1.2 × 10^−6^, FC = 1.09). No significant interactions were seen with mutations at TSSs and open-chromatin sites (**Supplementary Figure 6A**). *ARID1A* is a tumor suppressor encoding a member of the SWI/SNF chromatin remodeling complex that regulates chromatin accessibility at regulatory elements and is involved in the maintenance of genomic stability and DNA repair ^54^. *ARID1A* interacts with the topoisomerase TOP2A to facilitate its DNA binding and DNA decatenation, a process required for proper chromosome segregation in mitosis ^55^. The SWI/SNF complex is also involved in DNA double-strand break repair ^56^ and nucleotide excision repair ^57^, and loss of *ARID1A* impairs polymerase pausing ^58^. *ARID1A* is also one of the most frequently mutated genes in pancreatic adenocarcinoma ^59^ and in the PCAWG cohort, as 16/24 of tumors carried frameshift or stop-gain mutations that suggest loss of function. The inactivating *ARID1A* mutations may increase the mutation burden at TSSs as a result of impaired repair mechanisms and increased instability, and consequently, further contribute to transcriptional dysregulation.

In melanoma, driver mutations in *BRAF* were associated with mutational enrichment in CTCF binding sites (interaction *P* = 0.022) (**Figure 5C**). The 28 tumors with *BRAF* mutations were enriched in CTCF binding site mutations (*FDR* = 5.8 × 10^−5^, FC = 1.10), while 32 *BRAF*-wildtype tumors showed no enrichment (*FDR* = 0.31, FC = 1.03). BRAF serine/threonine kinase is a proto-oncogene and the activating V600E mutation defines a druggable subtype of melanoma ^60,61^. Previous work shows that ectopic expression of V600E-mutant *BRAF* in epithelial cell lines induces DNA double stranded breaks and the production of reactive oxygen species ^62^. In this cohort, 22/28 melanomas carried V600E substitutions and V600K substitutions occurred in three additional tumors. No significant interactions of *BRAF* mutations and localised mutagenesis were found for TSSs and open-chromatin sites (**Supplementary Figure 6B**). This analysis suggests that melanomas defined by *BRAF* driver mutations may have increased activity of a mutational process acting specifically on CTCF binding sites.

Amplifications of the 17q23.1 locus associated with an increased mutation frequency in CTCF binding sites in breast cancer (interaction *P* = 2.3 × 10^−4^) (**Figure 5D**). The 64 tumors with amplifications were enriched in mutations in CTCF binding sites (FDR = 1.1 × 10^−5^, FC = 1.11) while 127 tumors lacking the amplifications showed no enrichment (FDR = 0.91, FC = 1.00). This highest-ranking interaction was also confirmed in the pancreatic cancer cohort (P = 0.0058) where 17 tumors with amplifications showed significantly more mutations at CTCF binding sites (FDR = 1.7 × 10^−4^, FC = 1.18) compared to the 211 tumors lacking the amplifications (FDR = 0.042, FC = 1.04). Besides 17q23.1, we found 12 amplified loci in breast cancer with significant interactions with CTCF binding sites to potentially explain these effects. Also, chromosomal instability has been associated with mutagenesis of CTCF binding sites in gastrointestinal cancers ^28^. However, no significant interactions of high aneuploidy and CTCF binding site mutations for breast cancer were detected in our models. Discovery of the interaction in two cancer types leads to the speculation that the 17q23.1 locus is involved in localised mutagenesis through unknown mechanisms, however more work is needed to investigate this hypothesis in detail.

To further decipher the interactions of recurrent genomic amplifications and local mutagenesis, we determined the genes located in the amplified loci that responded transcriptionally to amplifications. Using a cancer-type specific analysis, we found 282 unique genes in 25 amplified regions that were significantly up-regulated in the tumors with these amplifications (Wilcoxon test, *FDR* < 0.05) (**Figure 5E**) (**Supplementary Table 1I**). mRNA abundance associations were identified for all three types of genomic sites and six cancer types. 13 known cancer genes were found (*RAD21, CDK12, ERBB2, RECQL4, CCND1, DEK, BCL11A, MYC, JAK2, PDCD1LG2, BRAF, EXT1, PTK6*).

In breast cancer, *RAD21* mRNA abundance was significantly higher in 8q23.3-amplified tumors compared to non-amplified tumors (*FDR* = 1.6 × 10^−7^) (**Figure 5F**), indicating that *RAD21* up-regulation is driven in part by the genomic amplification. The 8q23.3 amplification was associated with mutational enrichments in CTCF binding sites in breast cancer (*P* = 0.0089) (**Supplementary Figure 7A**). *RAD21* encodes a subunit of the cohesin complex that co-binds DNA with CTCF to orchestrate transcriptional insulation and chromatin architectural interactions ^48,49^, suggesting its role in CTCF-related mutagenesis.

As another example, *BRAF* was up-regulated in the subset of breast cancers with 7q34 amplifications (*FDR* = 0.0081) (**Figure 5G**). The amplification was associated with increased mutagenesis at TSSs in breast cancer (interaction *P* = 0.012) (**Supplementary Figure 7B**). The 7q34 amplification was also significantly associated with CTCF binding site mutations in esophageal cancer (interaction *P* = 0.034) (**Supplementary Figure 7B**), however no matching RNA-seq data was available to confirm the transcriptional upregulation of *BRAF*. Thus, our analysis associates *BRAF* amplifications and point mutations to localised mutagenesis in multiple cancer types.

In summary, this integrative analysis of mutational processes with recurrent driver mutations, copy number alterations and gene expression data provides a catalogue of somatic alterations that associate with localised mutational enrichments in gene-regulatory and chromatin architectural elements of the cancer genome. Since our evidence remains correlative, further computational and experimental approaches are needed to establish causal relationships. While a subset of these driver mutations and recurrent copy-number amplifications may be directly involved in mutagenesis and DNA repair, others may represent markers of tumor subtypes with specific exposures or endogenous factors. Deeper study of these associations may help decipher genetic mechanisms underlying mutational processes.

## Discussion

The cancer genome is molded by diverse mutational processes that continuously shape its broad megabase-scale features and the fine context of single nucleotides. Here we focused on the mutational processes of an intermediate scale that affect thousands of genomic elements, each spanning tens to hundreds of nucleotides. Such functional elements are widespread in the non-coding genome and play various roles in gene expression control and chromatin architecture. Using our novel computational framework, we characterised the mutational landscape of gene-regulatory and chromatin architectural elements in diverse cancer types and identified their putative functional and genetic determinants. We show that the ubiquitous mutational enrichment in TSSs is associated with high transcription and with biological processes involved in cellular housekeeping, but importantly with a variety of pathways and processes implicated in cancer. In contrast, transcriptionally silent genes showed no local deviations in mutation frequencies. On the other hand, open-chromatin regions were generally enriched in mutations and showed no association with transcript abundance. CTCF binding sites exhibited a particular pattern of site-specific mutagenesis: a small fraction of hypermutated sites defined by a core CTCF DNA motif and constitutive binding across tens of human cell types dominated over the majority of sites that lacked mutational enrichment. Lastly, an integrative analysis of site-specific mutation frequencies with recurrent alterations in cancer genomes allowed us to predict genetic drivers of localised mutational processes. The association of driver mutations in *ARID1A* and *BRAF* with enrichment of site-specific mutations corroborate earlier functional and mechanistic evidence of mutagenesis and DNA repair disruption, lending confidence to our other identified interactions with driver genes and genomic amplifications, and providing mechanistic hypotheses for future studies.

Open-chromatin regions were broadly enriched in mutations in most cancer types and the effect appeared to be independent of transcript abundance of predicted target genes. These findings contrast an earlier report that indicated decreased mutation rates in open-chromatin sites of cell lines at a similar resolution of the genome ^29^. The use of non-matched chromatin data of cell lines and fewer cancer genomes may explain the depletion signals observed earlier that may be confounded by tissue-specific properties of chromatin state and mutagenesis. Our analysis of the open-chromatin regions of matching cancer types and a larger set of whole cancer genomes likely provides a more accurate view of mutational processes. In our study, the pan-cancer analysis of open-chromatin sites also revealed only attenuated effects of total mutational enrichments and several mutational signatures, as well as indels appeared depleted in sites compared to flanking regions. Contrarily, interpreting individual cancer types with maps of cancer-specific open-chromatin sites revealed robust enrichments of mutations at open-chromatin sites. This highlights the importance of using matched genomic, epigenomic and transcriptomic profiles to study localised mutational processes.

We speculate that the local mutational enrichments in gene-regulatory and chromatin architectural sites represent a functional continuum of passenger and driver mutations. On the one hand, the vast majority of mutations enriched across a class of sites likely represent functionally neutral passenger mutations whose frequent occurrence is explained by differences in mutagenesis, DNA repair or carcinogen exposure. For example, previous studies of promoter-specific mutational enrichments in melanoma did not find broad associations of mutations and transcriptional deregulation of target genes ^63^, suggesting a lack of immediate functional impact. On the other hand, a small subset of genomic elements directly involved in the transcriptional or epigenetic control of hallmark cancer pathways may accumulate functional mutations either by chance or through the elevated activity of site-specific mutational processes. Those may show positive selection at an individual site or across a set of functionally-related sites. For example, while recent large-scale studies of whole cancer genomes have found relatively few individual non-coding regions with driver mutations ^4^, additional low-frequency candidate drivers were shown to converge onto the regulatory elements of cancer-related biological processes and protein interaction networks ^37^. Other recent studies have combined computational analyses and functional validation experiments to nominate new non-coding drivers in cis-regulatory modules and CTCF binding sites, and to associate these with transcriptional deregulation of cancer pathways ^34-36^. This suggests that our mechanistic understanding of oncogenesis, progression and metastasis pathways may be refined by analysing the non-coding genome. A better understanding of localised mutagenesis will help deconvolute the effects of mutational processes and positive selection and contribute to cancer driver discovery in the non-coding genome.

Our analysis has certain caveats and limitations. We analysed a broad catalogue of genomic elements that provides a limited representation of the heterogeneous dataset of tumor genomes.

We used several strategies to prioritise functionally active elements in a tissue-specific manner: TSSs were grouped by target gene transcription in the matching tumor samples, open-chromatin sites were selected from genome-wide profiles of cancers of matching types or related normal tissues, and CTCF binding sites were grouped by their binding conservation across a large panel of human cell types. To better address tumor heterogeneity, future analyses will benefit from detailed multi-omics cohorts where precisely matching genomic, transcriptomic and epigenomic profiles of individual tumors are available. Also, the current method is designed for genomic elements of uniform width and it is not directly applicable to elements of variable width, such as exons or non-coding RNAs. Our analysis suggests that different classes of gene-regulatory and architectural elements of the genome may be subject to localised mutational processes that have footprints of different sizes. Thus, it is recommended to evaluate the relevant input parameter of the method when analysing new classes of genomic elements. Our method is designed to quantify localised differences of mutation frequencies acting on an entire class of genomic elements with thousands to hundreds of thousands of genomic loci. It is not powered to evaluate a single genomic element as a potential cancer driver and alternative methods should be used for this purpose. However, we have adapted our method to evaluate TSS-specific mutation rates in gene sets of biological processes and pathways with hundreds to thousands of genes.

Our study opens new avenues for future developments. Integrative analysis of whole cancer genome sequences and rich clinical and pathological profiles of tumors ^17^ may highlight associations of clinical variables and localised mutagenesis and thus lead to the discovery of novel WGS-based biomarkers. Considering patient lifestyle information, environmental exposures and germline variation in the analysis may elucidate the impact of carcinogens and endogenous DNA repair deficiencies. Our catalogue of genetic associations provides hypotheses on mutational mechanisms that can be tested experimentally using genome editing and mutagenesis assays. Rare germline variants in the human population ^64^, *de novo* variants detected in genetic disorders ^21^ and the widespread somatic genome variation found in healthy tissues ^65^ provide further avenues to study mutational processes acting at functional non-coding elements. Our study provides a detailed annotation of localised mutational processes in whole genomes and enables future work to decipher the interplay of local mutagenesis and cancer driver mechanisms, molecular heterogeneity and genome evolution.

## Methods

### Regression models for Localised Mutations (RM2)

Local differences in mutation rates in functional genomic elements (*i*.*e*., sites) were evaluated using a negative binomial regression model we refer to as RM2. Single nucleotide variants (SNVs) and small insertions-deletions (indels) were analysed. The model simultaneously considers a collection of sites, such as regulatory elements that are commonly ∼10–1,000 bps in length and measured in ChIP-seq and related experimental assays in thousands to hundreds of thousands of genomic loci. Sites were uniformly redefined using their median coordinate and added sequences of fixed width upstream and downstream of the sites (*e*.*g*., ±25 bps or 50 bps around the midpoints of CTCF binding sites). Upstream and downstream flanking sequences of these sites were used as control regions to estimate expected mutation rates. Control regions were defined to be of equal width to sites such that the upstream and downstream regions combined were twice as wide as the sites initially. Next, site and flank sequences were pooled, duplicate sequences were removed, and overlapping coordinates of sites were removed from the flanking set such that every nucleotide and every mutation was considered as either as part of a site or a flanking control sequence but never both. The pooling and deduplication steps were used to account for groups of adjacent sites. To account for megabase-scale variation in mutation frequencies, we computed the total log-transformed mutation count for each site within its one-megabase window (i.e., ±0.5 Mbps around site midpoint). Based on this estimate, all sites were distributed into ten equal bins (*MbpRate*). The value of ten bins worked well in our benchmarks and captured variation in smaller and larger cohorts of individual cancer types. However, custom values of this parameter can be used. Mutation rates for sites and flanking sequences for each bin were defined separately and were distinguished by a binary cofactor (*isSite*). Sequence positions were counted separately by their trinucleotide context (*nPosits*) and expanded to three alternative nucleotides to account for the potential sequence space where such single nucleotide variants could occur (*nPosits*). The observed mutations in these contexts were also counted (*nMuts*) and a cofactor was used to add separate weights to different trinucleotide classes (*i*.*e*., reference trinucleotide and alternative nucleotide; *triNucMutClass*). Indels were counted under another entry in *triNucMutClass* such that all mutation counts were summed, and the entire genomic space was accounted for. An optional binary cofactor (*coFac*) was included to allow the consideration of genetic or clinical covariates of localised mutation rates. To evaluate the significance of localised mutation rates in sites compared to flanking control regions, we first constructed a null model that excluded the term *isSite*:

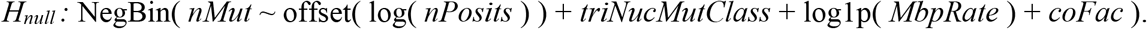

The main model representing the alternative hypothesis of a site-specific mutation rate was constructed as follows:

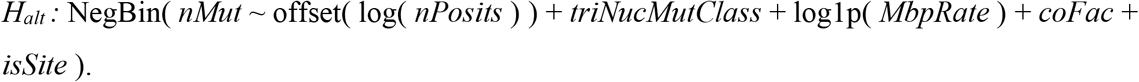

We extended our model to evaluate whether localised mutation rates differ between two subtypes of tumors, such as those defined by clinical annotations or genetic features using the term *coFac*. Trinucleotide sequence content, trinucleotide mutational signatures and megabase-scale covariations of mutation rates were computed separately for the two sets of tumors. To establish the associations of localised mutation rates and tumor subtypes (or presence of driver mutations), we added to the initial model the term *isSite:coFac* mapping the interaction of the tumor subtype and the cofactor distinguishing sites and flanking sequences, as follows:

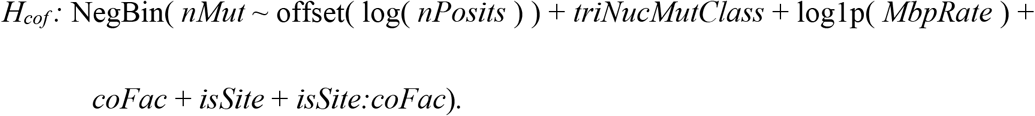

We used likelihood ratio tests to compare the models and evaluate the significance of localised mutation frequencies (*H*_*alt*_ *vs. H*_*null*_ to evaluate the term *isSite*). Chi-square *P*-values from the likelihood ratio tests were reported for each analysis. We also reported coefficient values of the term *isSite* to characterise enrichment or depletion of mutations at sites relative to flanking controls for positive and negative values, respectively. The interactions of driver mutations and mutation rates were evaluated using likelihood ratio tests that compared the models *H*_*alt*_ and *H*_*cof*_. Only the models with significant positive coefficients were reported. The expected mutation counts were derived from each model by 1000-fold sampling of mutation counts from the negative binomial distribution informed by the fitted probabilities and theta values derived from the regression models. Fold-change values were derived by dividing median observed and expected mutation counts, and confidence intervals were derived using the 2.5^th^ and 97.5^th^ percentiles of sampled values. Chi-square *P*-values from the models were corrected for multiple testing using the Benjamini-Hochberg procedure. As an exception, unadjusted P-values were used to quantify the interactions of local mutation frequencies and tumor subtype (*e*.*g*., presence of driver mutation) since these analyses were conditional of the significance of main site-specific effects. Besides modelling total mutations in sites and flanking sequences, we evaluated mutations of multiple subclasses, such as mutations stratified by transcriptional activity, COSMIC mutational signatures or DNA strands. Mutation subclass analysis was conducted as described above. The same megabase-scale mutation rates estimated for all mutations were used rather than those of specific subclasses. The method is available at https://github.com/reimandlab/RM2.

### Somatic mutations in whole cancer genomes

Somatic single nucleotide variants (SNVs) and short insertions-deletions (indels) in the genomes of 2,583 primary tumors were retrieved from the uniformly processed dataset of the Pan-cancer Analysis of Whole Genomes (PCAWG) project of the ICGC and TCGA ^3^. We used consensus variant calls mapped to the human genome version GRCh37 (hg19). We removed 69 hypermutated tumors with at least 90,000 mutations, resulting in 2,514 tumors and 24.7 million mutations. We also removed 33 tumors for which mutational signature predictions were not available in PCAWG. We analysed tumor genomes of the pooled pan-cancer cohort of multiple cancer types, and also 25 cohorts of specific cancer types with at least 25 samples in the PCAWG cohort. We excluded a small subset of tumors in which localised mutation rates were exceptionally strong even when analysing one tumor genome at a time (*FDR* < 0.001, RM2). The small subset of filtered tumors included 33 tumors (1.4% of the cohort) that provided 38 findings of elevated site-specific mutation frequencies of the three site classes. The outlier tumors showed overall higher mutation burden compared to the remaining pan-cancer cohort (mean 50,238 *vs*. 9,404 mutations). We performed this filtering because based on our initial analyses, we found that the individual contribution of these tumors to the overall analysis would have caused overestimates of mutation rates. The filtering led to a conservative analysis that only revealed associations shared across multiple tumor genomes. To enable this filtering, we performed tumor specific analyses for the three classes of sites (open-chromatin sites, CTCF binding sites, and TSSs). We analysed each cohort of a cancer type separately and grouped the mutations according to tumor ID, allowing the model to learn an expected background mutation rate in the respective cohort and then test each tumor genome separately for localised mutation rates. To perform this single-tumor analysis in smaller cohorts within the PCAWG dataset (<25 tumors of a given type), we created a meta-cohort by pooling these smaller cohorts. After filtering hypermutated tumors, tumors without PCAWG signatures, and tumors with exceptionally strong signals of localised mutations, we derived a conservative final set of 2,419 genomes of 35 cancer types with 22.7 million mutations including 1.61 million indels. To evaluate the performance of our model, we also processed a dataset of simulated variant calls for the same set of tumors derived from the PCAWG project (*i*.*e*., the Broad dataset) ^4^. The true and simulated datasets were compared for method benchmarking (see below). In addition to evaluating total mutations, several classes of mutations were analysed separately. Mutations were mapped to C and T nucleotides and grouped by reference and alternative nucleotides (C>[A,G,T], T>[A,C,G]). Mutations were also classified as located either on the Watson (w) strand if the original reference nucleotide was C or T, or the Crick (c) strand if the original reference nucleotide was A or G. We also classified mutations by the trinucleotide signatures of single base substitutions (SBS) that were derived earlier using the SigProfiler software in the PCAWG project ^14^. We assigned each mutation to its most probable signature in the given patient tumor based on its trinucleotide context. For model evaluation, mutations were also annotated for the simulated dataset.

### Chromatin architectural and gene-regulatory genomic elements

We performed a systematic analysis of three classes of genomic elements: DNA-binding sites of CTCF (CCCTC-binding factor) detected in multiple human cell lines, transcription start sites (TSS) of protein-coding genes, and open-chromatin sites (ATAC-seq sites and DNAse-seq sites) detected in human primary tumors or related normal tissues. CTCF binding sites were retrieved from the ENCODE project ^39^. Sites observed in only one cell line were removed for the majority of the study, resulting in 119,464 sites across 70 cell lines. As an exception, the full set of 162,209 CTCF binding sites in ENCODE was used for the analysis of CTCF binding sites that considered site conservation across cell lines (Figure 4), as discussed below. TSS loci of protein-coding genes were retrieved from Ensembl Biomart (GRCh37) and filtered based on location of standard chromosomes (1-22, X, Y), resulting in 37,309 TSSs of 18,710 protein-coding genes. Open-chromatin sites were collected from ATAC-seq profiles of a TCGA study ^40^ and DNAse-seq profiles of the Roadmap Epigenomics project ^41^. We used tissue-specific open-chromatin profiles for every cancer type, first by matching cancer types of the PCAWG project and the TCGA project (17/25 cohorts), and second, by selecting the closest normal tissues or cell lines of the Epigenomics Roadmap project for cancer types for which the cancer-specific open-chromatin profiles were not available (8/25 cohorts) (**Supplementary Table 1A**). The pan-cancer analysis considered pan-cancer open-chromatin sites of the TCGA ATAC-seq study. To better compare the mutation rates of TSSs and open-chromatin sites, we excluded the open-chromatin sites that overlapped with the TSSs or CTCF binding sites. Sites that were originally mapped to the GRCh38 reference genome were re-mapped to GRCh37 using the LiftOver method. Throughout the study, the three classes of sites were normalised to uniform width based on median coordinates. CTCF binding sites were defined using 50 bps (±25 bps) windows around the midpoint of sites. Midpoints of TSS loci were defined in the Ensembl database and we used a 200 bps (±100 bps) window around the TSSs. Open-chromatin sites were also defined using a 200 bps (±100 bps) window around site midpoints. We systematically explored various values of the site width parameters and the final selection was based on the strength of signal and consistency (**Supplementary Figure 2A**).

### Defining target genes of TSSs and open-chromatin sites

TSS and open-chromatin sites were analysed in groups based on the mRNA abundance of target genes in matching tumors. TSS target genes were retrieved from the Ensembl database and target genes of open-chromatin sites were predicted using two custom strategies. First, direct target genes of open-chromatin sites were defined based on their location in gene promoters, 5’ and 3’ UTRs, and exons (open-chromatin sites directly overlapping TSSs were excluded, as specified above). Second, a promoter-capture Hi-C dataset from a recent multi-tissue study ^46^ was used to identify long-range chromatin interactions of open-chromatin sites and target genes. We selected high-confidence interactions with frequency above 10 that were observed in at least 5/28 cell types to define putatively constitutive interactions of open-chromatin sites and target genes.

### Grouping TSSs and open-chromatin sites by tissue-specific mRNA abundance

We used mRNA abundance in matching tumors as a proxy of site functionality of TSSs and open-chromatin sites. To group sites by mRNA abundance, we used the uniformly processed PCAWG RNA-seq dataset ^45^ (RPKM-UQ) that covered ∼50% of the whole cancer genome cohort. We limited the mRNA analysis to the tumors in the WGS dataset and excluded non-coding genes. Cancer type specific analyses were carried out in 20 cohorts of cancer types for which at least 15 tumor samples with mRNA and WGS data were available, and also the pooled pan-cancer group as the 21^st^ cohort. We discarded genes with duplicated HGNC symbols and genes for which TSS or open-chromatin sites were not mapped. Together, this resulted in mRNA measurements for 20,042 protein-coding genes in 1,267 tumor transcriptomes. Next, for each cancer type, we grouped all the genes into five bins based on their median mRNA abundance and analysed the bins using RM2 separately for TSSs and open-chromatin sites. For the pan-cancer analysis, we binned genes using median mRNA abundance in the entire RNA-seq dataset.

### Grouping CTCF binding sites by the conservation of CTCF binding across human cell lines

To analyse CTCF binding sites by their tissue and cell type specificity, we grouped all 162,209 CTCF binding sites of the ENCODE dataset into five equally sized bins based on the number of cell lines where a binding event of CTCF to the sites was detected. To interpret these CTCF binding sites, we used information on the three-dimensional genome and CTCF sequence motifs. First, we retrieved chromatin loops in eight cell lines from a Hi-C study ^48^, used a ±1,000 bps window around loop anchor midpoints to define narrower versions of loop anchors, and counted the number of CTCF binding sites in each bin overlapping these loop anchors. We used a Fisher’s exact test to evaluate the enrichment of CTCF binding sites at loop anchors among the CTCF binding sites with constitutive activity across cell types (i.e., the 5^th^ bin of CTCF sites). Second, using the motif MA0139.1 of the JASPAR database ^66^, we detected the presence of the core CTCF motif in CTCF binding sites using the FIMO method ^67^ and selected significant motif instances (*FDR* < 0.01). Enrichment of CTCF motifs in the 5^th^ bin of sites was identified similarly.

### Analysis of localised mutation frequencies in gene-regulatory and chromatin architectural elements

First, we evaluated the localised mutation rates in CTCF binding sites, TSSs and cancer-specific open-chromatin sites for the pan-cancer cohort and all cohorts of selected cancer types. Total mutations and mutations grouped by COSMIC signatures and reference/alternative allele were analysed. Indel mutations were analysed as part of total mutations and also as a separate group. Results were adjusted for multiple testing using the Benjamini-Hochberg false discovery rate procedure and filtered (*FDR* < 0.05). We also analysed the simulated variant call set using the same pipeline and found no significant results, as expected (*FDR* < 0.05). Results of the systematic analysis were visualised as a dot plot. *FDR* values in the main dot plot were capped at 10^−32^ for visualisation purposes. To visualise localised mutation rates, all sites were pooled, aligned using median coordinates and trimmed to uniform lengths. Coordinates were transformed relative to site midpoint. Upstream and downstream flanking sequences of equal length were also considered. Local regression (LOESS) curves with the span parameter of 33% were used to visualise a smoothened mutation frequency in sites relative to flanking sequences.

### Associating mutations in TSSs and open-chromatin sites with transcript abundance

To study the associations of mRNA abundance and mutations in TSSs and open-chromatin sites, we grouped the sites by mRNA abundance of genes in matching tumor types, and then performed the RM2 analysis separately for each bin and cancer type. Since RM2 is better powered to discover localised mutation rate differences in larger groups of sites, we used down-sampling of TSSs and open-chromatin sites in the mRNA-defined gene bins to provide better comparisons. In each bin and cancer type, we randomly sampled 4,500 sites for RM2 analysis and repeated this procedure over 100 iterations and selected the RM2 result with the median P-value for reporting. The number for down-sampling was chosen as the closest smallest common value for bin sizes for TSSs and open-chromatin sites. The results were adjusted for multiple testing using the Benjamini-Hochberg false discovery rate procedure and filtered (*FDR* < 0.05).

### Identifying pathways with frequent mutations in transcription start sites

We asked whether the localised changes in mutation frequencies of TSSs affected specific biological processes and pathways. We repurposed the RM2 model to analyse TSSs of gene sets corresponding to biological processes of Gene Ontology ^68^ and molecular pathways of the Reactome database ^69^. Gene sets were derived from the g:Profiler ^70^ web server (March 3^rd^, 2020) and subsequently filtered to include 1,871 gene sets with 100 to 1,000 genes while smaller and larger gene sets were excluded. Results were corrected for multiple testing using the Benjamini-Hochberg FDR procedure separately for every cancer type and filtered for statistical significance (*FDR* < 0.05). The pathways with significantly higher TSS-specific mutation frequencies were visualised as an enrichment map ^71^ in Cytoscape and major biological themes were manually curated as described earlier ^47^. Nodes in the enrichment map were painted to reflect cancer types where these pathway enrichments were detected according to the custom color scheme of the PCAWG project.

### Associating elevated mutation frequencies in CTCF binding sites with constitutive CTCF binding and core sequence motifs

To study the functional associations of localised mutation frequencies at CTCF binding sites, we first analysed five groups of CTCF binding sites defined based on the counts of cell lines where the sites were observed. These bins, ranging from sites of single cell lines to sites constitutively active in all or most cell lines, were analysed in all cohorts of individual cancer types and the pan-cancer cohort. Findings were corrected for multiple testing and filtered to select significant findings (*FDR* < 0.05). To perform the motif-based analysis, we first split sites into two groups based a significant match to the core CTCF motif using FIMO ^67^ (FDR < 0.01). Then, we repeated the binning of sites by shared activity, as described above, to ensure all bins of sites within each group (with or without motif) were of equal size. The ten bins of CTCF sites were also analysed using RM2. Lastly, to perform a tissue-specific analysis of matching cancer types and cell lines, we selected CTCF binding sites for two cancer cell lines in the ENCODE dataset: MCF-7, a breast cancer cell line, and HepG2, a liver cancer cell line. The CTCF binding sites found in at least these two cell lines were also split into ten bins based on conservation of CTCF binding and presence of the core sequence motif, and then analysed with RM2 using mutations of the PCAWG dataset for the Breast-AdenoCa cohort (with MCF-7) and Liver-HCC cohort (with HepG2).

### Associating elevated mutation frequencies in sites with driver mutations and recurrent copy-number alterations

We tested whether the localised changes in mutation frequencies in CTCF binding sites, TSSs and open-chromatin sites were associated with driver mutations (*i*.*e*., SNVs, indels) or recurrent copy-number alterations (CNAs). First we collected a high-confidence set of driver mutations and CNAs in the PCAWG cohort. Driver mutations in exons of protein-coding genes were predicted for each selected cancer type using the ActiveDriverWGS method ^34^. For driver analysis, we used the PCAWG variant calls after filtering tumors as described above, corrected the results for multiple testing using the Benjamini-Hochberg FDR procedure and selected significant driver genes (*FDR* < 0.05). FDR correction was conducted separately for each cancer type across the pooled set of protein-coding and non-coding genes. Tumors with and without SNVs or indels in predicted driver genes were used for localised mutation rate analysis. Predictions of recurrent CNAs were derived from the pan-cancer dataset of GISTIC2 calls of the PCAWG project ^53^. All lesions at 95% confidence scores were considered and amplifications and deletions were analysed separately. High-confidence CNA events were used (GISTIC2 score = 2). Tumors with and without CNAs in the recurrently altered regions, as defined by GISTIC2, were used for localised mutation rate analysis. Two additional chromosomal features were used. First, we grouped the tumors by presence or absence of Whole Genome Duplication (WGD) as determined in the PCAWG project ^53^. Second, we computed the Percent Genome Altered (PGA) metric for every tumor as a proxy of aneuploidy. PGA was computed as the percentage of the autosomal genome affected by CNA segments whose total copy number deviated from the global reference (two copies for non-WGD genomes; four copies for WGD genomes). PGA metrics were median-dichotomised separately for every cancer type, resulting in two subgroups of tumors: the PGA-high group and the PGA-low group corresponding to above-median and below median CNA burden of tumors. Following the construction of these genetic features (*i*.*e*., driver gene mutations, recurrent CNAs, WGD, PGA-high), we filtered overly frequent (more than 2/3 of the cohort) and infrequent features (less than 15 tumors) to improve the power of the RM2 analysis. For each cancer type, the relevant features were then analysed for localised elevations in mutation frequencies in the three categories of genomic elements (open-chromatin sites, CTCF binding sites, TSSs). The binary co-factor in RM2 was used to indicate the presence or absence of the given feature in the given tumor genome. We first computed the significance of site-specific localised mutation rates given the presence or absence of the feature. To validate this joint model, subgroups of tumors with and without the defining feature were also analysed separately using RM2. All combined RM2 results concerning individual features, cancer types and genomic sites were then adjusted for multiple testing correction using the Benjamini-Hochberg FDR procedure and significant results were selected (*FDR* < 0.05). We filtered the results to only include positive and significant interactions of genetic features and elevated localised mutation frequencies (interaction *P* < 0.05, main and interaction coefficients > 0) and compared the FDR values, fold-change values and visualisations of localised mutation frequencies to validate these findings.

### Associating mutagenesis-related copy number amplifications with mRNA abundance

To evaluate the functional role of recurrent copy number amplifications associated with increased mutation frequencies in the gene-regulatory and chromatin architectural sites, we studied the genes located in the amplified regions and retrieved from the PCAWG GISTIC2 dataset. We compared the mRNA abundance levels of the genes between groups of tumors defined by the presence or absence of the amplifications, using matching RNA-seq data available in PCAWG ^45^. Genes with low mRNA abundance were removed from the analysis (median FPKM-UQ < 1). mRNA abundance levels of genes in amplified and non-amplified (*i*.*e*., balanced and deleted) tumors were compared using one-sided non-parametric Wilcoxon tests, assuming that an increase in mRNA abundance would match the underlying copy number amplification. Results were adjusted for multiple testing correction using the Benjamini-Hochberg FDR procedure and significant results were selected (*FDR* < 0.05). We ranked the resulting genes based on the total significance of CNA/mRNA associations across all cancer types and site types by summing the negative log10-transformed significant FDR-values of each gene. Known cancer genes of the COSMIC Cancer Gene Census database ^72^ (v91, downloaded 14.05.2020) were highlighted.

### Method benchmarking and power analysis

We evaluated the performance of our method and statistical power using simulated variant calls, different parametrizations and down-sampling of input datasets. First, to evaluate method calibration and false-positive rates, we performed a systematic analysis of open-chromatin sites, TSSs, and CTCF binding sites in a comparable set of simulated variant calls from PCAWG. This simulated variant set was derived earlier from the same set of tumor genomes using trinucleotide-informed shuffling of mutations ^4^. Simulated variant calls were analysed similarly to true variant calls for total mutation counts, reference and alternative nucleotide combinations and predicted mutational signatures. Results from RM2 were adjusted for multiple testing using the Benjamini-Hochberg FDR procedure separately for results derived from true and simulated variant calls. As expected, the simulated variant calls revealed no statistically significant results of localised mutation rates in any cancer type, site type or mutation subset (*FDR* < 0.05). We then visualised the distribution of log-transformed *P*-values derived from true and simulated variant calls using quantile-quantile (QQ) plots and found that highly significant *P*-values were detected in true datasets while the *P*-values derived from simulated variant calls were uniformly distributed as expected. These analyses show that our model is well-calibrated and is not subject to inflated false-positive findings. Second, to evaluate the statistical power of RM2, we performed a systematic down-sampling analysis by randomly selecting subsets of sites and tumors for localised mutation rate analysis. We focused on the PCAWG liver hepatocellular carcinoma (Liver-HCC) cohort of 300 samples and CTCF sites. A series of down-sampling configurations were used (2000, 5000, …, 100,000 sites sampled; 25, 50, …, 250, 300 genomes sampled), as well as the full datasets of sites and genomes. Each configuration was tested 100 times with different random subsets of data. As a power analysis, we evaluated the fraction of runs that revealed a significant enrichment of somatic mutations at CTCF sites (*P* < 0.05) and the median *P*-value of these 100 runs. Third, we evaluated the parameter values of RM2 that determine the genomic width of each site and the control regions of upstream and downstream flanking sequences. As expected, site-specific mutation rates were consistently identified at multiple values of the width parameter for each class of site (open-chromatin sites, CTCF binding sites, and TSSs), indicating robustness of our analysis to different parameter values. However, different site classes showed preferences towards shorter sites (CTCF binding sites: 20-100 bps) or longer sites (open-chromatin sites and TSS: 200-800 bps), likely due to different underlying mutational processes. For the entire study, the optimal genomic size of every site class was selected based on the strongest effect size and significance across multiple cancer types. The value of 50 bps (±25 bps) was selected for CTCF sites. For open-chromatin sites and TSSs, we selected the common site width of 200 bps (±100 bps) that showed strong effects in both TSSs and open-chromatin sites, to increase comparability of the two classes. We evaluated the effect of grouping sites by their megabase-scale mutation frequencies (*i*.*e*., the MbpRate covariate of RM2). We repeated the systematic analysis using a simplified RM2 model that excluded the MbpRate covariate and compared the resulting P-values and FDR values derived from the original and simplified models. The original model (including MbpRate) outperformed the simplified model (excluding MbpRate) by assigning stronger significance to findings in the dataset of true mutations and reduced significance to findings in the simulated dataset, indicating that accounting for megabase-scale mutation frequency as covariate improves sensitivity and specificity of the model.

## Acknowledgments

This work was supported by a Project Grant from the Canadian Institutes of Health Research (CIHR) to J.R., an Operating Grant from the Cancer Research Society to J.R., and the Investigator Award to J.R. from the Ontario Institute for Cancer Research (OICR). C.A.L. was partially supported by a Graduate Student Fellowship from the Department of Medical Biophysics, University of Toronto. Funding to OICR is provided by the Government of Ontario.

## Author contributions

J.R., C.A.L. and D.A.R. designed and implemented the method. J.R. and C.A.L. analysed the data. J.R. wrote the manuscript with significant input from C.A.L. and D.A.R. J.R. conceived and supervised the project. All authors reviewed and edited the manuscript and approved the final version.

