## Supplementary Figures for "Functional and genetic determinants of mutation rate variability in regulatory elements of cancer genomes"

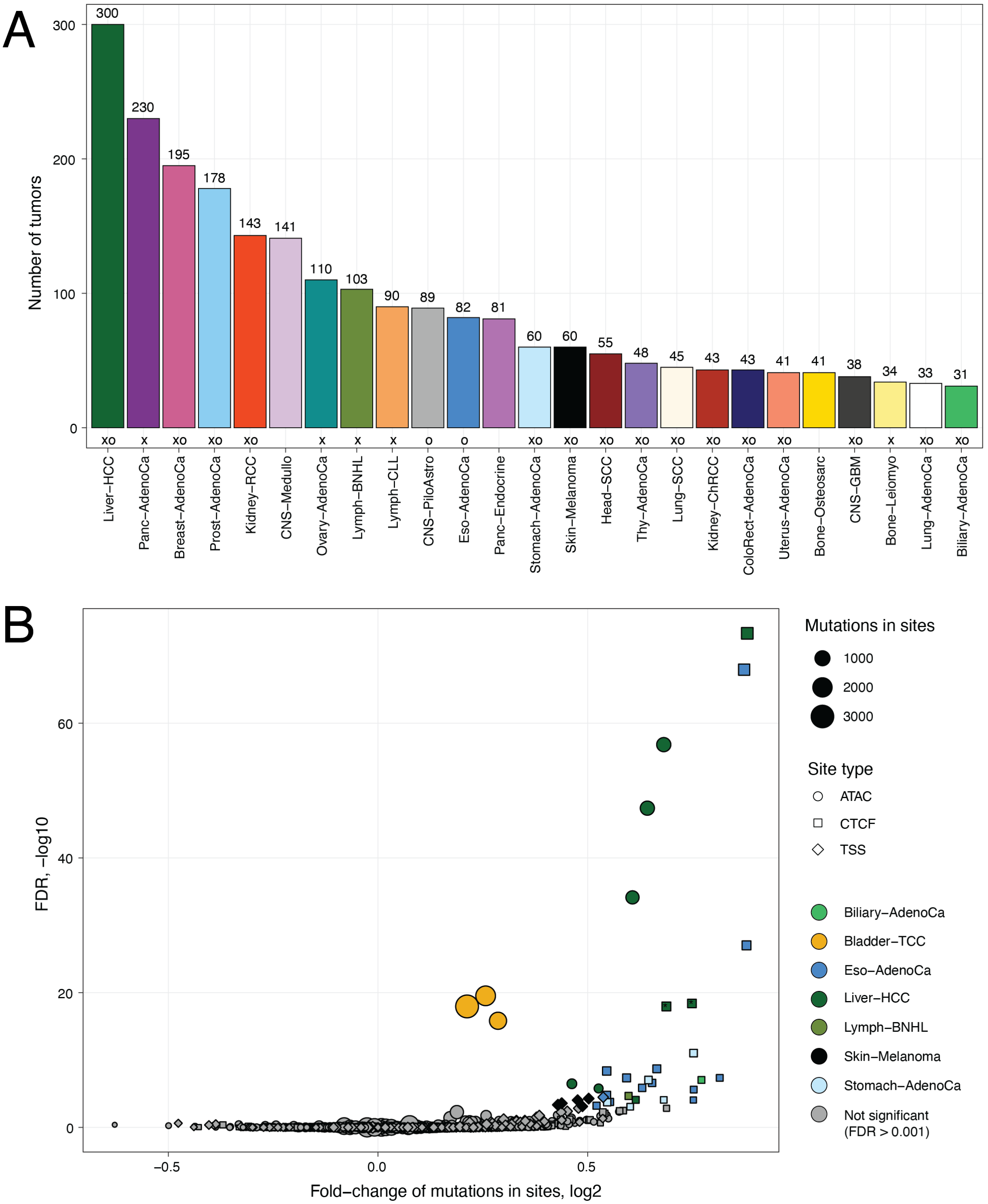


**Supplementary Figure 1. Dataset of whole cancer genomes used in the study. A.** The PCAWG pan-cancer dataset of 2,419 whole cancer genomes with 25 cohorts with at least 25 tumors was used for the analysis. We analysed the pan-cancer cohort of all tumors and the 25 cohorts separately. The ten cancer types with fewer samples are not shown. Symbols below bars denote matching datasets (X – RNA-seq of matching tumors in PCAWG; O – ATAC-seq of matching tumor types in TCGA). **B.** Volcano plot of outliers shows 38 cases (33 tumors) with highly significant site-specific mutation frequencies that were excluded to obtain a more conservative analysis ($FDR < 0.001$; colored dots). 33 tumors correspond to 38 cases since a few tumors were found as outliers in more than one site type.

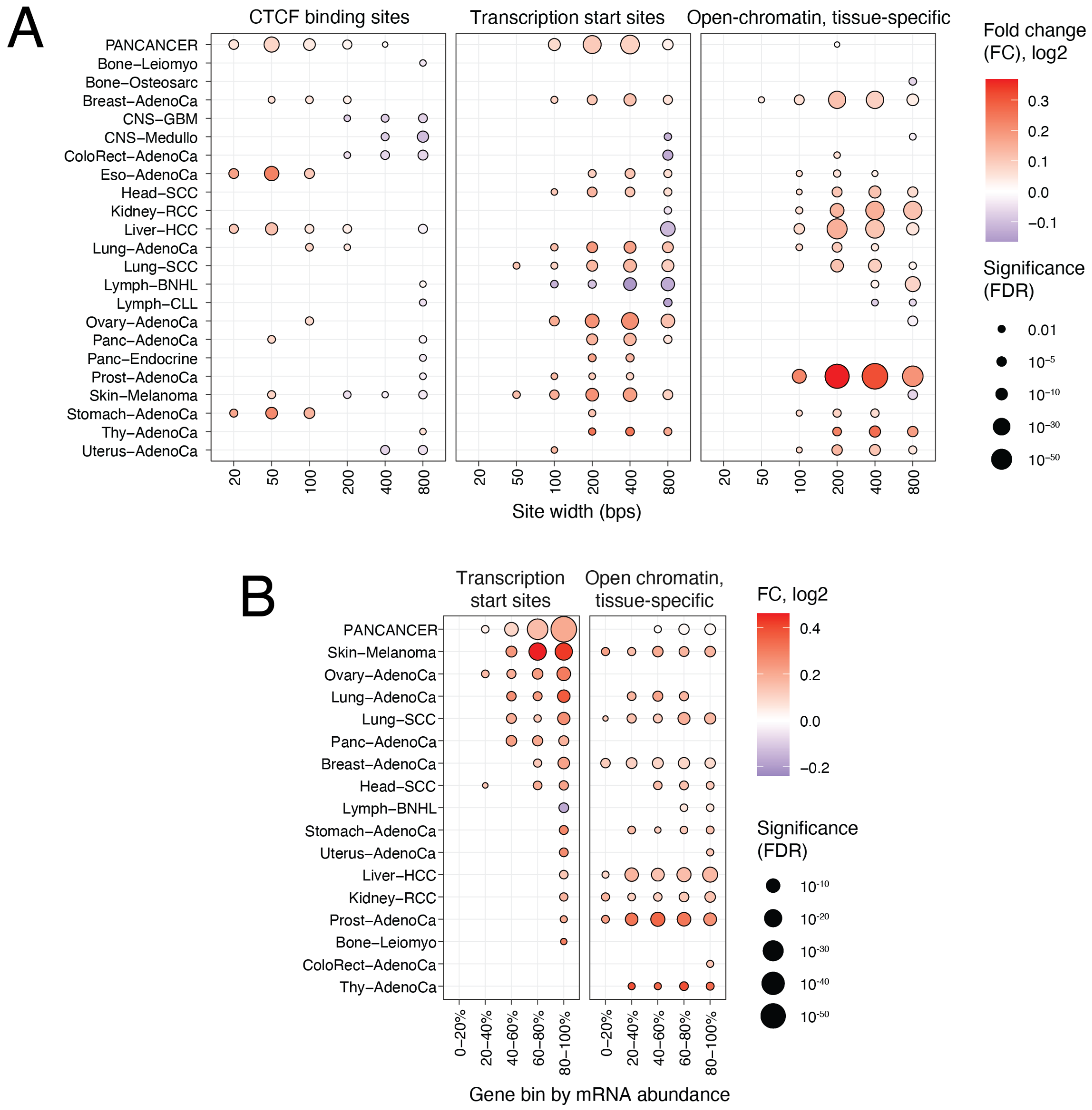


**Supplementary Figure 2A. Evaluating the impact of the size width parameter on RM2 analysis**. To evaluate the robustness of our method, we used different sizes of sites (*i.e.*, sequences of fixed lengths around site midpoints) to detect localised changes in mutation frequencies. The site width parameter also determines the size of flanking control sequences around sites. Mutational enrichments at CTCF sites were generally detected at smaller sites compared to open-chromatin sites and TSSs. Sites of 50 bps (CTCF binding sites) and 200 bps (TSS, open-chromatin) were used. Site width (X-axis) shows the number of bases considered for each site and its upstream and downstream flanks.

**Supplementary Figure 2B. Analysis of all TSSs and open-chromatin sites grouped by mRNA abundance of target genes.** Five genes bins of equal size were compiled for every cancer type based on median mRNA abundance in matching RNA-seq data. This supplementary analysis uses all sites per bin to evaluate local mutation frequencies, in contrast to the analysis in Figure 3 that uses down-sampling to include smaller random sets of sites to provide an unbiased comparison of TSSs and open-chromatin sites. Here, open-chromatin sites show stronger enrichments of mutations in multiple cancer types, consistent with our overall analysis of open-chromatin sites. The mutational enrichment signals in additional cancer types are partially explained by improved power enabled by larger sets of sites.

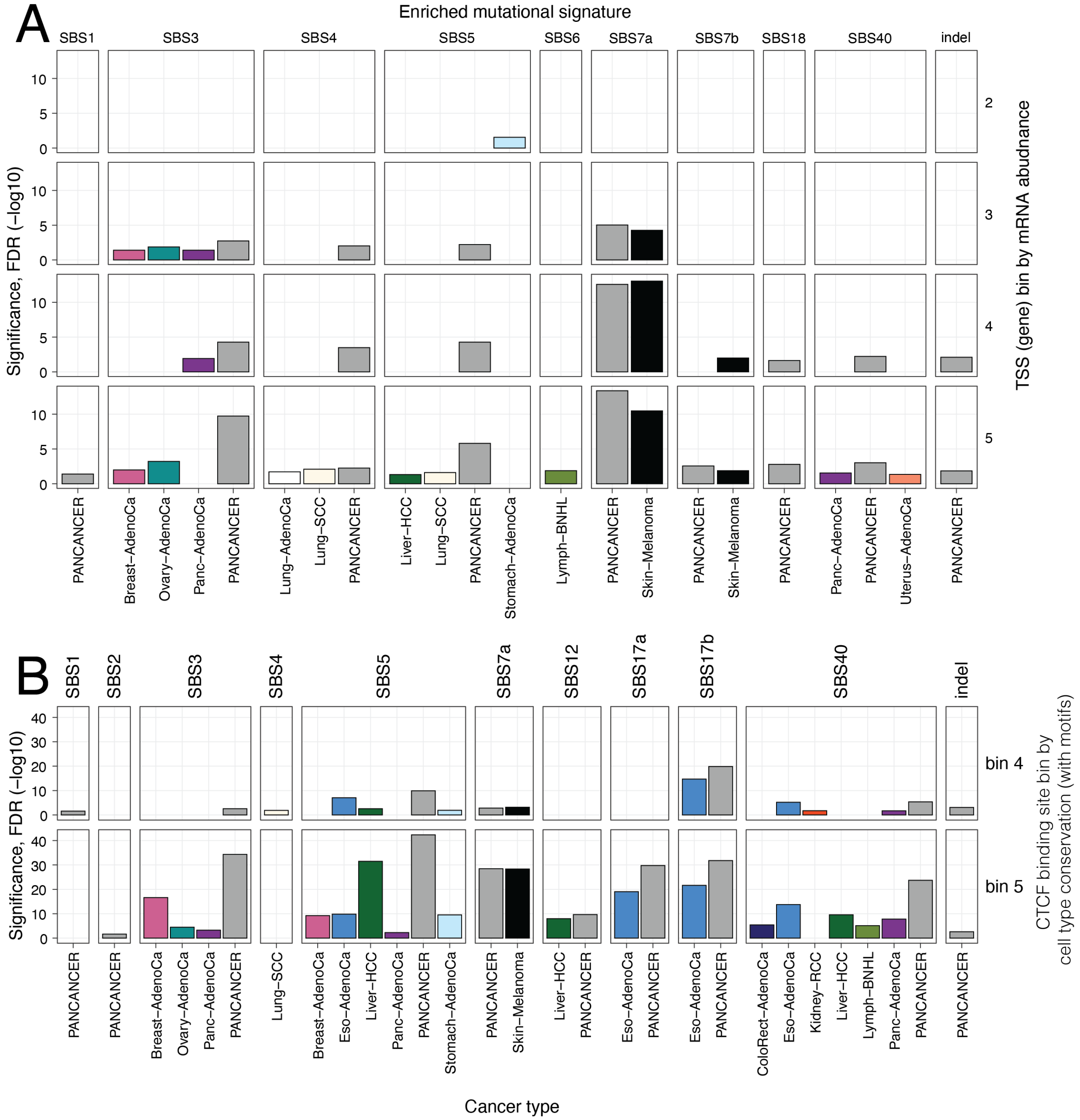


**Supplementary Figure 3. Enrichments of specific mutational signatures localised in active transcription start sites (TSS) and CTCF binding sites**. **A.** Mutational signature analysis of TSSs grouped by mRNA abundance of target genes (corresponding to Figure 3). The fifth bin of sites represents the top 20% of genes with the highest median mRNA abundance in matching cancer samples. **B.** Mutational signature analysis of CTCF binding sites grouped by the activity of sites across the panel of cell lines in ENCODE (corresponding to Figure 4). The fifth bin of sites shows CTCF binding activity in most or all ENCODE cell lines. In both analyses, sites comprising the fifth bin, corresponding to the most active sites, are the most enriched in a variety of mutational signatures.

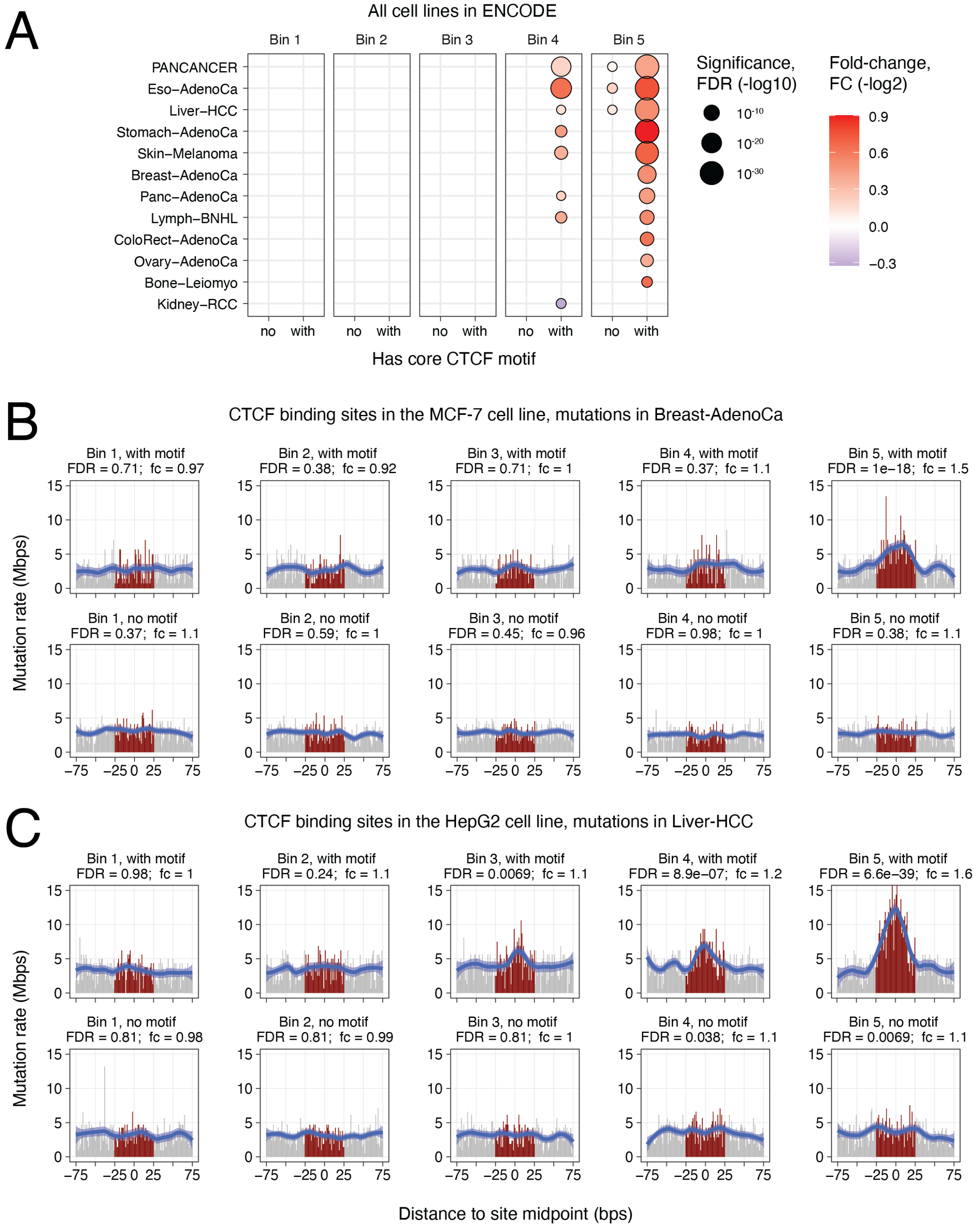


**Supplementary Figure 4. Comparison of localised mutation frequencies in CTCF binding sites that include or lack a core CTCF DNA-binding sequence motif. A.** Mutational enrichments of the five bins of CTCF binding sites based on the activity in 70 cell lines, grouped by the presence or absence of a CTCF sequence motif. Increased mutation frequencies affect constitutively active CTCF binding sites that include a high-confidence motif. In contrast, the other site bins are not enriched in mutations. **B-C.** Analysis of tissue-specific CTCF binding sites with mutations of the matching cancer type confirms motif-associated mutational enrichments in constitutively active, motif-including CTCF binding sites. **B.** Enrichment of liver cancer mutations (Liver-HCC) in CTCF binding sites detected in at least the HepG2 liver cancer cell line. **C.** Enrichment of breast cancer mutations (Breast-AdenoCa) in CTCF binding sites detected in at least the MCF-7 breast cancer cell line.

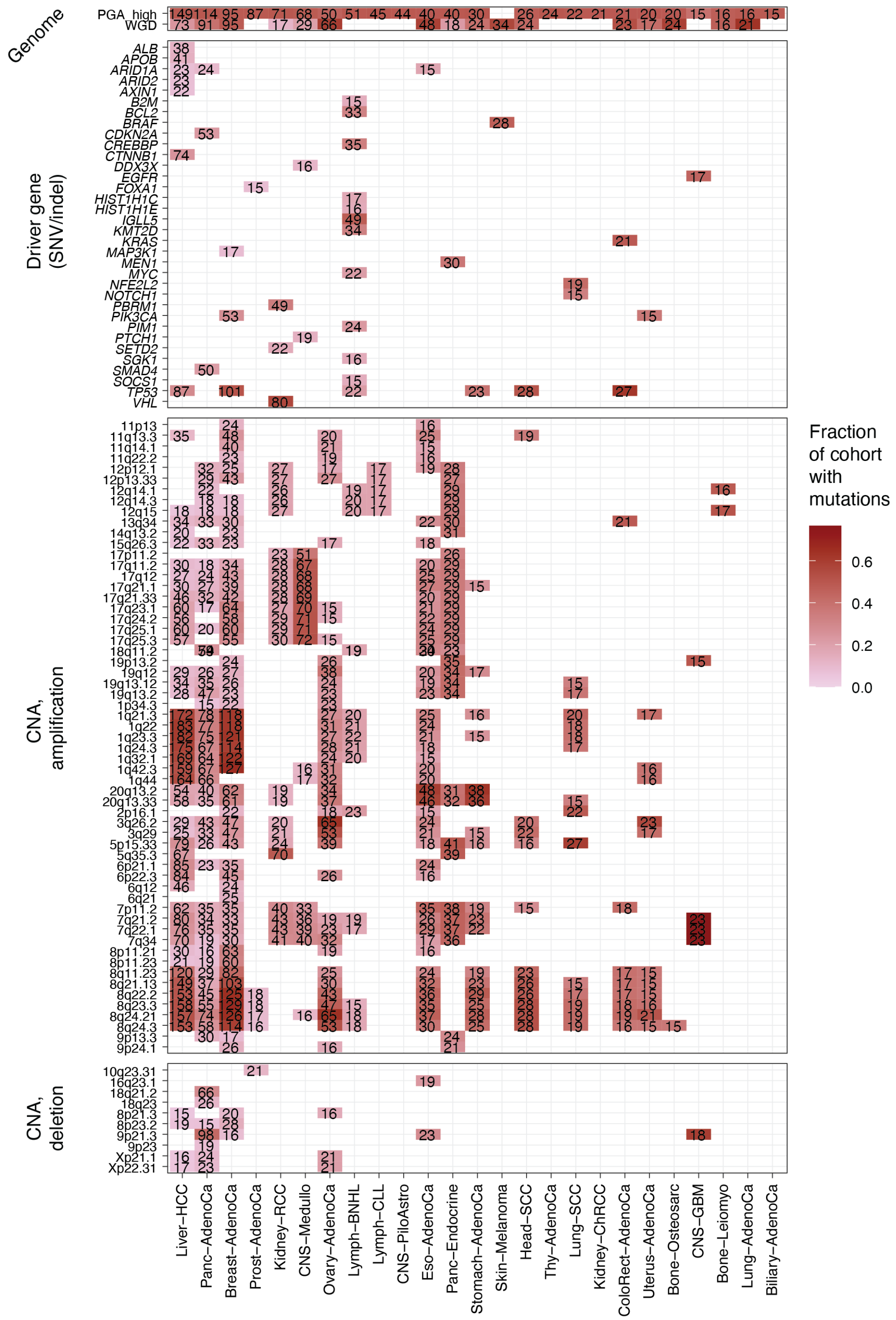

**Supplementary Figure 5. Driver gene mutations and recurrent copy-number alterations (CNA) considered for analysis of mutation frequencies.** Grid plot shows the driver genes and CNAs analysed for associations with mutation frequencies. Counts of tumors with mutations are shown in cells. For each cancer type, driver mutations and CNAs in at least 15 tumors and in no more than 2/3 of the cohort were used. Driver genes were predicted using ActiveDriverWGS. Recurrent amplifications and deletions were derived from the GISTIC2 analysis of the PCAWG project. Two genome-wide features of whole-genome duplication (WGD) and percent genome altered (PGA) were also included.

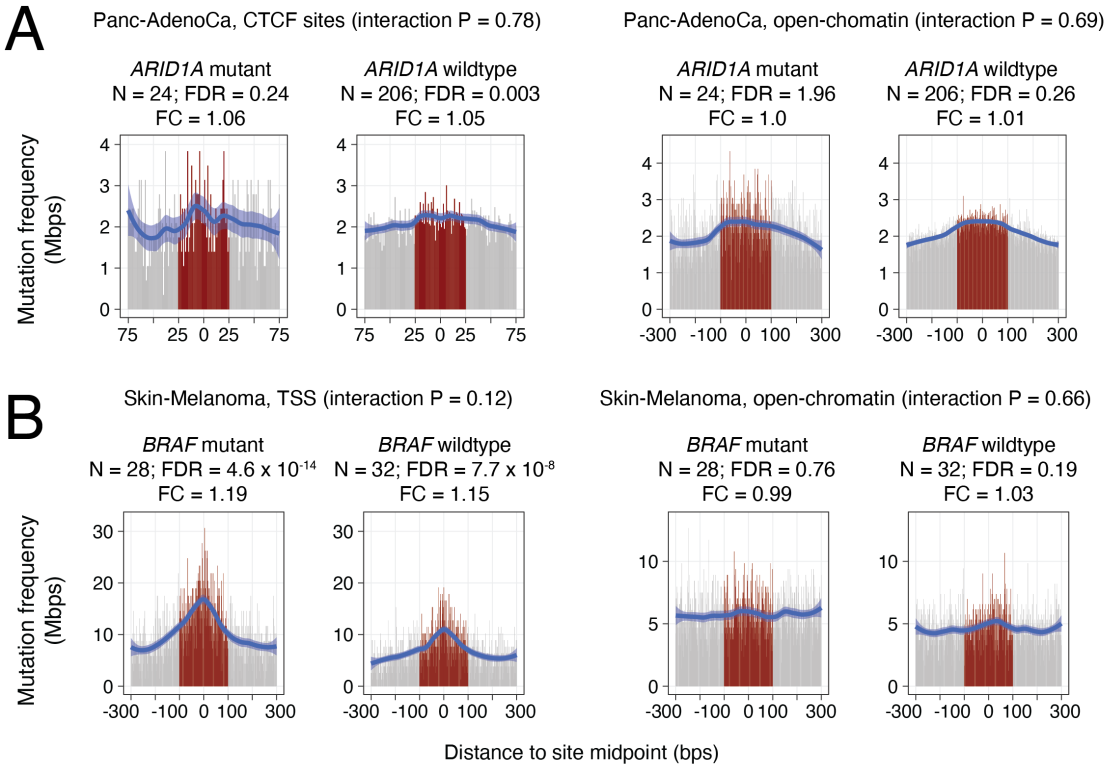


**Supplementary Figure 6. Interactions of local mutation frequencies and predicted drivers. A.** Control comparisons for *ARID1A* in pancreatic cancer. No significant interactions of mutations and local mutation frequencies in CTCF binding sites (left) and open-chromatin sites (right) were found. **B.** Control comparisons for *BRAF* in melanoma. No significant interactions of mutations and local mutation frequencies in TSSs (left) and open-chromatin sites (right) were found. Driver-wildtype (left) and driver-mutant tumors (right) are compared within each group. Number of tumors, FDR and fold-change values and interaction P-values from RM2 are shown.

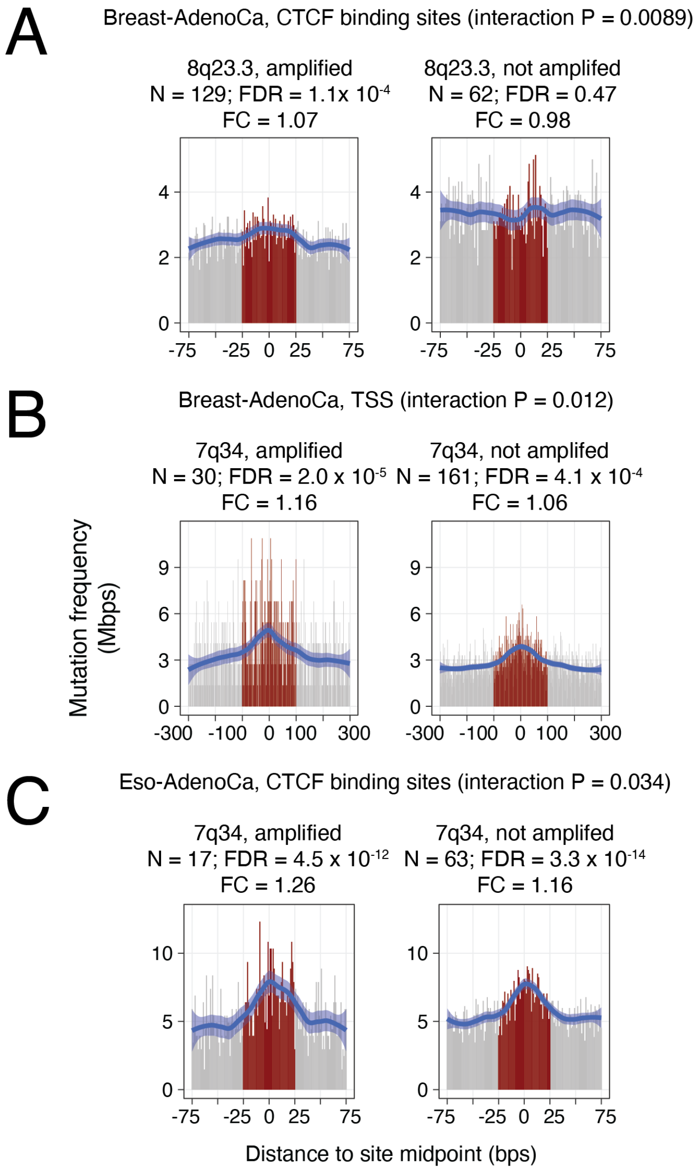


**Supplementary Figure 7. Additional interactions with copy-number amplifications and increased mutation frequencies in genomic elements.** Tumors with and without specific recurrent amplifications are compared (left *vs.* right). **A.** Amplifications in the 8q23.3 locus encoding *RAD21* are associated with an increased mutation frequency of CTCF binding sites in breast cancer. **B.** Amplifications in the 7q34 locus encoding *BRAF* are associated with an increased mutation frequency of TSSs in breast cancer. **C.** The 7q34 amplification also associates with frequent mutations in CTCF binding sites in esophageal cancer. Number of tumors, FDR and fold-change values and interaction P-values from RM2 are shown.
